# The Hippo pathway transcriptional co-activator YAP is involved in head regeneration and bud development in *Hydra*

**DOI:** 10.1101/2021.03.24.436861

**Authors:** Manu Unni, Puli Chandramouli Reddy, Sanjeev Galande

## Abstract

The Hippo signaling pathway has been shown to be involved in the regulation of cellular identity, cell/tissue size maintenance and mechanotransduction. The Hippo pathway consists of a kinase cascade which determines the nucleo-cytoplasmic localization of YAP in the cell. YAP is the effector protein in the Hippo pathway which acts as a transcriptional cofactor for TEAD. Phosphorylation of YAP upon activation of the Hippo pathway prevents it from entering the nucleus and hence abrogates its function in transcription of target genes. In Cnidaria, the information on the regulatory roles of the Hippo pathway is virtually lacking. Here, we report for the first time the existence of a complete set of Hippo pathway core components in *Hydra*. By studying their phylogeny and domain organization, we report evolutionary conservation of the components of the Hippo pathway. Protein modelling suggested conservation of YAP-TEAD interaction in *Hydra*. We also characterized the expression pattern of the homologs of *yap, hippo, mob* and *sav* in *Hydra* using whole mount RNA in situ hybridization and report their possible role in stem cell maintenance. Immunofluorescence assay revealed that *Hvul*_YAP expressing cells occur in clusters in the body column and are excluded in the terminally differentiated regions. The YAP expressing cells are recruited early during head regeneration and budding implicating the Hippo pathway in early response to injury or establishment of oral fate. These cells exhibit a non-clustered existence at the site of regeneration and budding, indicating the involvement of a new population of YAP expressing cells during oral fate specification. Collectively, we posit that the Hippo pathway is an important signaling system in *Hydra*, its components are ubiquitously expressed in the *Hydra* body column, and may play crucial role in *Hydra* oral fate specification.

## Introduction

The ability of the cells to come together and act in a coordinated fashion helped organisms evolve from solitary single-cell based forms to extremely complex and specialized multicellular organisms. Signaling pathways which allow cell-cell communication in a highly specific and spatio-temporal manner enabled the advent of multicellularity. This is evident while studying ontogenesis. Considering the sheer number of cell-types present in a complex organism such as humans, it comes as a surprise that the development of an organism from a zygote to a fully formed adult is controlled by a complex interplay of merely 10 main classes of signaling pathways (Perrimon et al., 2012). These include- Notch, Wnt, Hedgehog, TGF**β**/BMP, Receptor-tyrosine kinase (RTK), Hippo, NF-***κ***B, JAK-STAT, JNK & Nuclear receptor signaling pathway family.

Multicellularity arose about 400-1000 million years ago on earth (Butterfield, 2000) independently in at least 16 different eukaryotic lineages which led to complex multicellular taxa like metazoa, Fungi and Embryophyta (Butterfield, 2000;Brunet and King, 2017). Considering the bilaterians as the most complex and diverse multicellular clade, a basic ‘Developmental Toolkit’ required for generation, organization and maintenance of multicellular structures can be assessed. The origin of these developmental tools which include transcription factors, signaling pathways, cell adhesion and cell polarity related genes can be traced back to basal metazoans (Tweedt and Erwin, 2015). A detailed analysis of these development toolkits, body plan and differential germ layers and a diverse cell-type system indicates Cnidarians are arguably the first phylum to evolve and exhibit features which underlie the traits commonly seen in Bilateria. Cnidarians exhibit an oral-aboral body axis polarity with a diploblastic germ layer organization. These germ layers in cnidarians have been reported to form myoepithelial cells, nerve-net of sensory/ganglion neuronal cells, gastric cells, germline cells and cnidocytes which are the defining feature of the phylum. Studies in the past few decades have shown clearly that these primitive organisms display highly complex developmental programs and toolkits which are commonly found in the bilaterians.

Among the cnidarians, *Hydra* is the best-characterized model. *Hydra* is a freshwater polyp known to exhibit tremendous regenerating potential with a capability to regenerate even from reaggregated cells of dissociated polyps (Gierer et al., 1972). It has been a classical model for developmental and regeneration biology for more than two centuries and has contributed immensely towards the understanding of morphogen mediated processes and understanding various cell signaling pathways (Reddy et al., 2019b). Among the ten developmentally important signaling pathways as discussed above, seven of them have been shown to be functional according to the studies on *Hydra*. Many components of the Wnt signaling have been reported in *Hydra* and their role has been established to be important in the regulation of head organizer activity (Hobmayer et al., 2000). Many components of Notch signaling are present in *Hydra* and have been reported to be important in the boundary formation in tissues (Sprinzak et al., 2010; Münder et al., 2013). The TGFβ superfamily of signaling pathway has also been reported to be crucial for *Hydra* developmental signaling such as during tentacle formation, foot formation, symmetry breaking (Reinhardt et al., 2004; Rentzsch et al., 2007; Watanabe et al., 2014). Members of the RTK family of signaling pathways - VEGF, FGF and Ephrin have been shown to be crucial for regeneration in *Hydra* (Tischer et al., 2013; Krishnapati and Ghaskadbi, 2014). NF-***κ***B has been reported to be important for early regenerative time points in *Hydra* (Franzenburg et al., 2012; Wenger et al., 2014). While their role is presently thought to be innate immunity/inflammation-related, its direct developmental regulation is yet to be established. JNK in *Hydra* has been found to be crucial in nematocyte differentiation and regulation of TLR-signaling (Philipp et al., 2005; Franzenburg et al., 2012). Among the nuclear receptor family of signaling pathways, Retinoblastoma gene has been found to be expressed in almost all cell types in *Hydra* but its specific role has not been deciphered (Schenkelaars et al., 2018). Another nuclear receptor protein, NR3E has been found to be expressed in *Hydra* and is predicted to respond to the parasterol A, a cnidarian A-ring aromatic steroid (Khalturin et al., 2018). Among the three remaining developmentally important signaling pathways yet to be reported in *Hydra* are- Hedgehog, JAK-STAT and Hippo signaling.

Hippo pathway has emerged as a major player for the orchestration of spatio-temporal regulation of cell differentiation, proliferation, tissue size control, and apoptosis. These capabilities enable the Hippo pathway to be important in the regulation of morphogenesis and tissue or organ regeneration. It was first described and reported in *Drosophila* while screening for tumour suppressor genes in 1995 (Xu et al., 1995). However, it was only in 2005 when Yorkie (Yki), a transcription co-activator, was linked to Hippo signaling, and the importance of the Hippo pathway in regulating transcriptional landscape was truly realized (Huang et al., 2005). Yes-associated protein, also known as YAP, is a highly conserved mammalian homolog of the *Drosophila* Yki. The Hippo core components are kinases which phosphorylates YAP through a cascade, which represses its transcriptional activity by preventing its nuclear transportation and hence its interaction with transcription factors like TEAD (Fulford et al., 2018). Upon phosphorylation, YAP is sequestered in the cytoplasm through 14-3-3 interaction or undergo ubiquitination for its degradation. The core components of Hippo characterized in *Drosophila* consists of Ser/Thr kinases- Hippo (Hpo) and Warts (Wts); and their adapter proteins- Salvador (Sav) and Mats. In mammalians, the equivalent set of factors is named as-Mst, Lats, Sav and Mob, respectively. At the cellular and molecular levels, the Hippo pathway and its functions are highly conserved between invertebrates and vertebrates.

There has been a paucity of literature to date about Hippo signaling in basal metazoans. The role of Hippo signaling in highly regenerative organisms like *Hydra* is unknown. A recent study in another cnidarian reported that *Clytia hemispherica* has all the core components of Hippo pathway and CheYki has cell proliferation regulatory function. Hence, it is pertinent to characterize the homologs of core Hippo pathway components in *Hydra* to understand their role in *Hydra* regeneration and cell proliferation and differentiation. A recent study has reported the presence of core components of the Hippo pathway in *Nematostella* (Hilman and Gat, 2011), suggesting that this pathway has fairly conserved ancient origin during the evolution of multicellular organisms. Here, using a combination of bioinformatic analysis and molecular cloning, we report the existence of a complete set of core Hippo pathway components in *Hydra.* Using domain analysis and 3D protein modelling, we show that these homologs have a conserved domain and motif architecture indicating a possible conserved interactive signaling network. Whole-mount *in situ* hybridization (WISH) analysis revealed that these genes are expressed across the body column with a few gene-specific variations. Adapting CheYki specific antibody for immunofluorescence assay of *Hydra* YAP, we show that nuclear localized YAP occurs as clustered cells across the body column with no expression in the regions which are terminally differentiated. We show that the YAP expressing cells are recruited to the regenerating tip and early buds. Further, we report the existence of a separate non-clustered nuclear localized YAP expressing cell population at the hypostomal region which may be involved in oral fate specification and maintenance.

## Materials and Methods

### Animal culture

Clonal culture of *Hydra vulgaris* Ind-Pune (Reddy et al., 2011) was maintained in *Hydra* medium by following standard methods at 18±1°C (Horibata et al., 2004). Polyps were fed daily with freshly hatched Artemia nauplii larvae and washed 6–8 h after feeding. For regeneration experiment, *Hydra* polyps starved for 24 hrs were decapitated just below the tentacle base and allowed to regenerate till 0, 1, 2, 4 and 8 hours post amputation (hpa). The polyps were then fixed and processed for immunofluorescence assay. For budding experiment, *Hydra* polyps starved for 24 hrs were collected at different stages of bud development. These were then fixed and processed for immunofluorescence assay. The different stages of budding were identified and labelled as reported previously (Otto and Campbell, 1977).

### Identification of Hippo pathway homologs in *Hydra*

*Hydra magnipapillata* genome draft comprising 82.5% of 1.05 Gbp sequenced genome available as Refseq was initially used for identifying Hippo Pathway core components (Chapman et al., 2010). This assembly turned out to be incomplete and we were unable to fish out any homologs. An in-house transcriptome assembly generated in the Galande laboratory (Reddy et al., 2019a) was therefore used for the present study. To further improve the assembly, the in-house transcriptome was merged with the NCBI RefSeq to generate a hybrid assembly. The hybrid assembly was found to be 99.6 % complete as compared to 95.7% exhibited by NCBI RefSeq (Reddy et al., 2019a). Using the stand-alone NCBI BLAST program, hits of homologs of Hippo pathway core components were identified (Madden, 2013). To confirm the hits, Reverse BLAST was performed. Finding a hit of a homolog in different phyla or species would confirm the homolog status. To further confirm, the amino acid sequences of these homologs were searched in HMMER (Hmmer, RRID:SCR_005305) for affirmation based on the hits returned (Potter et al., 2018). Once the homologs were identified, further analyzed for domain organization by SMART (SMART, RRID:SCR_005026) (Letunic et al., 2002). After manual evaluation of the domain organization, the domain architecture was constructed to scale using DOG 2.0 software (Ren et al., 2009).

### Molecular phylogenetic trees

Sequences from different representative phyla were collected based on protein BLAST searches using Human YAP sequence as query. The collected sequences were aligned using MUSCLE (Edgar, 2004). The alignment was trimmed using automated trimAl programme (Capella-Gutiérrez et al., 2009). This alignment was subjected for phylogenetic analysis using FastTree 2 to generate an approximately maximum likelihood (ML) tree (Price et al., 2010). This method was selected after testing PhyML and RaxML as FastTree 2 has given better confidence on branching points and this could be due to highly divergence nature of the sequences. This pipeline was implemented in online platform NGphylogeny.fr (Lemoine et al., 2018). Here, LG substitution model was used with Felsenstein’s phylogenetic bootstrap with a value of 1000 (Lemoine et al., 2019). Phylogenetic tree was visualized by using iTOL webserver (Letunic and Bork, 2019). Tree was rooted using *Amphimedon queenslandica* YAP-like sequence as an outgroup. The domain organization analysis and visualization were carried out using DoMosaics software (Moore et al., 2014) using embedded HMMER3 tools (Mistry et al., 2013) and Pfam data. The sequences details were provided in Supplementary Table 1.

For the analysis of the rest of the Hippo pathway components, alignments and molecular phylogenetic trees of the protein sequences were carried out using MEGA 6.0 software (Tamura et al., 2013). MUSCLE algorithm was used for amino acid sequence alignment (Edgar, 2004). The alignment was graphically represented using Jalview (Waterhouse et al., 2009).

### Cloning of Hippo pathway homologs from *Hydra*

Total RNA was extracted from *Hydra* polyps starved for 48 hrs and cDNA was synthesized from total RNA using Improm-II reverse transcriptase system (Promega™) according to the manufacturer’s instructions. Hippo pathway genes were amplified by polymerase chain reaction using Pfu DNA polymerase with the following primers:

*Hvul_yap*_forward:5’ATGGATATGAATTCTACGCAACGGC3’,
reverse: 5’CTACAACCAAGTCATATATGCATTAGGC3’;
*Hvul_tead*_forward:5’ATGGCGGAAAACTGTCGAGATCC3’,
reverse: 5’TCAGTCTCTGACTAATTTAAATATGTGGT3’;
*Hvul_hpo*_forward:5’ATGTCTCGCAGTTTGAAGAAGTTGAG3’,
reverse: 5’TTAAAAATTTGCTTGCCTGCGTT3’;
*Hvul_mob*_forward:5’ATGAGTTTCCTGTTTGGCTCCA3’,
reverse: 5’TTATTTATTAATTAACTTATCCATAAGTTC3’;
*Hvul_lats*_ forward:5’ATGGCAGCTAATAATCTTTTTAGTAG3’
reverse: 5’TCATACAAAAACAGGCAACTTGC3’;
*Hvul_sav*_forward:5’ATGTTTAAGAAAAAAGATATTATCAAAACA3’,
reverse: 5’TTAAACATGAGTTTTTTTAAAAGAAATACT3’

#### The PCR conditions

Initial denaturation at 94°C for 5 min, followed by 30 cycles of denaturation at 94°C for 30 sec, annealing at the respective annealing temperatures (Ta) for 45 sec and extension at 72°C for 45 sec with the final extension 72°C for 5 minutes. The PCR amplified products were gel eluted using Mini elute kit (Qiagen), followed by A-tailing reaction using KapaTaq enzyme and cloned in pGemT-Easy vector system (Promega™) or TOPO TA cloning vector as per the manufacturer’s instructions. The recombinant plasmids were sequenced using sequencing primers and the nucleotide sequences of cloned genes were deposited at NCBI Genbank (*Hvul_yap*- MW650883; *Hvul_tead*- MW650884; *Hvul_hpo*- MW650879; *Hvul_mob*- MW650880; *Hvul_lats*- MW650881 and *Hvul_sav*- MW650882).

### Whole mount in situ hybridization

Digoxigenin-labelled sense and antisense RNA probes were prepared by in vitro transcriptions using recombinant plasmids of target genes made as mentioned above (Roche Life Science) and used for in situ hybridization. Whole mount in situ hybridization was performed on the polyps as described by Martinez et. al., (1997) with the following changes (Martinez et al., 1997). The animals were relaxed for 2 min in 2% urethane. Treatment with proteinase-K was performed for an optimum of 15 min and heat-inactivation of the endogenous alkaline phosphatases was done at 70°C for 15 min in 1X SSC. Digoxigenin labelled RNA probes at a concentration of 200-600 ng/ml of the probe was used for hybridization at 59°C. The post-hybridization washes were performed using 1X SSC-HS gradients. After staining with BM-purple AP substrate for 30 min-1 hr at room temperature, the animals were mounted in 80% glycerol for imaging. Imaging was carried out using 10 X DIC objective lens with Axio Imager Z1 (Zeiss).

### Cryosectioning of WISH stained *Hydra* samples

The stained polyps were rehydrated to PBS gradually through PBS: methanol gradient (25%, 50%, 75%, and 100 % wash each for 10 mins). These polyps were then shifted to a 30 % sucrose solution by gradually taking it through 10 % and 20 % for 30 mins each. The polyps were left in 30 % sucrose overnight. These polyps were then embedded in 10 % PVP (polyvinyl pyrrolidone) by making cubes of PVP (1 × 1 × 2 cm^3^) made from aluminum foil cast. The embedded polyps were then sectioned (25 μm thick) using Leica CM1950 – Cryostat. The sectioned ribbons were then collected on a glass slide and covered and sealed under a coverslip. The sectioned were then photographed under ZEISS Axio Zoom V16 apotome microscope.

### Analysis of expression of Hippo pathway components from single-cell transcriptome profile

t-SNE plots and gene expression plots of Hippo pathway components were generated and extracted from the Single Cell Portal (https://portals.broadinstitute.org/single_cell/study/SCP260/stem-cell-differentiation-trajectories-in-Hydra-resolved-at-single-cell-resolution). In order to use the Single Cell Portal, gene IDs of Hippo pathway components were acquired through a BLAST search in the Juliano aepLRv2 nucleotide database via *Hydra* 2.0 Genome Project Portal (https://research.nhgri.nih.gov/Hydra/sequenceserver/). To determine the clusters of cells that express individual Hippo pathway components, differential gene expression was analyzed using edgeR, which is a tool to analyze RNA-seq data using the trimmed mean of M-values (TMM) method. Differential gene expression was calculated as fold change.

### Immunofluorescence staining

A recently published paper reported the presence of Yorkie (YAP/Yki) in *Clytia hemispherica* and producing polyclonal antibody specific to CheYki in rabbit against the peptide FNRRTTWDDPRKAHS (Coste et al., 2016). This antibody along with the pre-immune serum was kindly gifted by Dr Michaël Manuel (Sorbonne Universités, Université Pierre et Marie Curie (UPMC), Institut de Biologie Paris-Seine (IBPS) CNRS). The antibody was validated by immunofluorescence analysis.

Immunofluorescence assay was performed as per the protocol is given in Takaku et. al., 2014 (Takaku et al., 2014). *Hydra* polyps were starved at least for one day before fixation. Animals were relaxed in 2% urethane for 1-2 min and fixed in 4% paraformaldehyde (in 1XPBS) overnight at 4°C or 1 hr at RT. 1:100 concentration of primary antibody was used. 1:100 concentration of Invitrogen Alexa-conjugated secondary antibodies was used. Invitrogen Alexa 488 conjugated Phalloidin for staining. DAPI was used for nuclear staining. These samples were then imaged on ZEISS Axio Zoom V16 (for regeneration and whole animal images) or Andor Dragonfly Spinning Disc (for budding *Hydra*) microscopes.

### Modelling

The 2.8 A° crystal structure of the human YAP-TEAD complex deposited on PDB (4RE1) was as a reference for modelling the TEAD binding domain and YAP binding domains of *Hydra* YAP-TEAD complex (Zhou et al., 2015). Modeller software (Webb and Sali, 2016) was used to build five optimum models based on 4RE1 in the multi-model mode. Among the five models, the model with minimum DOPE assessment score and maximum GA341 assessment score was chosen for the final analysis. The model was then visualized in CHIMERA for analysis, superpositioning and annotation (Pettersen et al., 2004). The non-covalent bond analysis was performed using Biovia Discovery Studio Visualizer (BIOVIA, 2017). The Binding energy calculations were performed using PRODIGY web server (Xue et al., 2016).

## Results

### Characterization and phylogenetic analysis of *Hydra* Hippo pathway genes

Core Hippo pathway homologs – *hippo/mst, mob, lats, sav, yap and tead* were identified from the in-house *Hydra* transcriptome using NCBI stand-alone BLAST (Reddy et al., 2019a). In mammals, *hippo, mob, lats* and *yap* have 2 paralogs each while *tead* has 4 paralogs. *Sav*, on the other hand, has no reported paralogs. The occurrence of these paralogs has been attributed to whole-genome duplication events correlated to certain fish species (Chen et al., 2019). Therefore, any species evolved earlier than fishes do not contain paralogs as reported for Hippo pathway genes. Conforming to these reports, *Hydra* consists of only one gene coding for each of the core Hippo pathway components. The *Hydra* Hippo pathway homologs were labelled as- *Hvul_hpo, Hvul_mob, Hvul_lats, Hvul_sav, Hvul_yap* and *Hvul_tead*. The presence of these homologs in *Hydra* was confirmed by obtaining the corresponding amplicons from *Hydra* cDNA (Supplementary Figure 1). Upon determining the nucleotide percent identity with other reported model organisms used for studying the Hippo pathway, we find that *Hydra* had a higher percent identity with humans than *Drosophila* (Figure 1 B). *Hvul_hpo* shows about 60 % identity with humans and 56 % identity with *Drosophila*. *Hvul_sav* is comparatively less conserved with a 24 % identity with human and 22 % identity with *Drosophila*. *Hvul_mob* is highly conserved across the animal phyla with about 84 % identity with humans and 83 % identity with *Drosophila*. *Hvul_lats* shares 41.6 % identity with humans and 42 % identity with *Drosophila*. *Hvul_yap* shows 34.6 % identity with humans and 34 % identity with *Drosophila*. *Hvul_tead* exhibits about 65 % identity with humans and 59 % identity with *Drosophila*.

**Figure 1:**
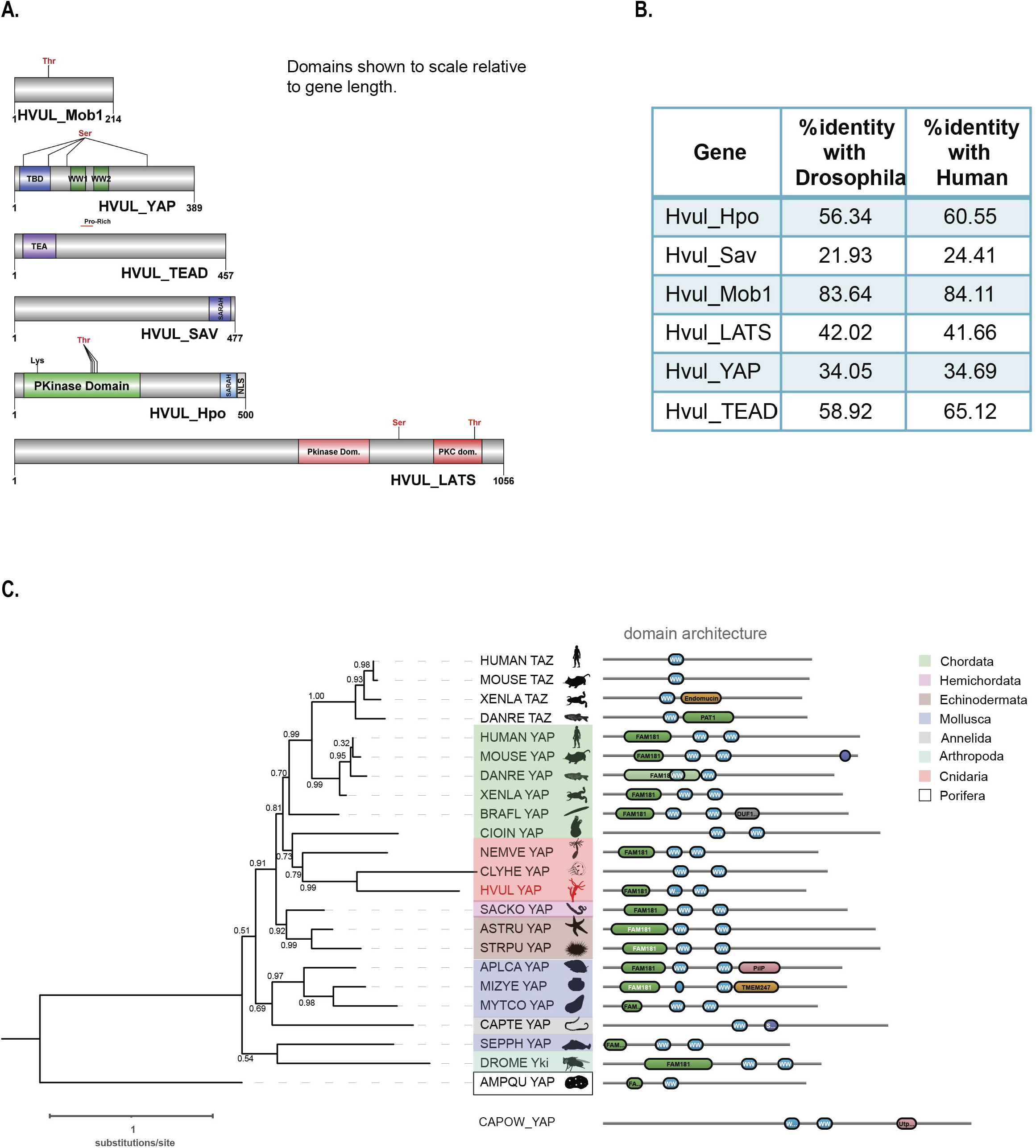
Identification of Hippo pathway homologs in *Hydra*. **A.** Domain architecture of the homologs as visualized using DOG 2.0. **B.** Depicts the percent identity of *Hydra* homolog with *Drosophila* and Human. **C.** Phylogenetic tree and domain organization of YAP homologs across the animal phyla. The phylogenetic analysis was carried out on NGphylogeny.fr webserver and the tree was generated using FastTree 2 method. Here, the phylogenetic tree was rooted at Amphimedon queenslandica YAP-like sequence (AMPQU). Domain organization analysis was carried out using DoMosaics software. Branch support values are displayed at the branching points. Different phyla are highlighted with distinct colours. *Hydra* YAP homologue (*HVUL* YAP) is highlighted in red colour font. A UNIPROT style abbreviations for organism names are used. Sequence details are provided in the Supplementary Table 1.

An earlier study performed the phylogenetic analysis of YAP homologues found in selective phyla (Hilman and Gat, 2011). However, this analysis did not cover majority of invertebrate phyla such as Annelida, Mollusca and Echinodermata. This could be due to lack of reliable data for the identification of the YAP homologues. Here, we have combined the phylogenetic analysis with predicted domain architecture. We used a YAP-like sequence found in *Amphimedon queenslandica* as an outgroup for rooting the tree. Additionally, a protein sequence with BLAST similarity from a unicellular Eukaryote (C*apsaspora owczarzaki*) was used for domain organization comparison. In this analysis, we observed that *Hydra* homologue of YAP exhibits a strong affinity to the chordate counterparts rather than non-chordate homologues (Figure 1C). An interesting observation after inclusion of multiple invertebrate phyla in the analysis is that they are highly diverged compared to the Cnidarian and Chordata species. This can be interpreted based on the weak branch support values (Figure 1C). Additionally, a molluscan homologue, *Sepia pharaonic* (SEPPH_YAP) showed more similarity with *Drosophila* Yki and S*accoglossus kowalevskii* homologue (SACKO_YAP) showed more similarity with Echinodermata homologues (Figure 1C). Domain organization analysis has led to identification of variability in the N-terminal homology domain (FAM181). This region contains TEAD binding domain (TBD). Surprisingly, in *Capitella teleta* (Annelida), *Clytia hemisphaerica* (Cnidaria) and *Ciona intestinalis* (Chordata) the FAM181 domain could not be detected (Figure 1C). This could be due to the higher sequence divergence in this region.

A detailed domain analysis using SMART website for the amino acid sequence of Hippo pathway homologs revealed a highly conserved domain organization of the proteins analyzed which indicates a fully functional pathway consisting of these core components (Figure 1 A). *Hvul*_HPO domain analysis revealed conserved N-terminal Protein kinase domain (PKinase Domain) and a C-terminal SARAH (Salvador-RASSF-Hippo) domain. The presence of these domains indicates the conserved regulation of activation of *Hvul*_HPO kinase activity (Glantschnig et al., 2002;Praskova et al., 2004;Boggiano et al., 2011). The *Hvul*_SAV also can be seen to have conserved the SARAH domain required for orchestrating the reported scaffolding activity (Yin et al., 2013). *Hvul*_LATS domain architecture indicates conservation of the hydrophobic motif (Motif: AFYEFTFRHFFDDGG) (a 40% hydrophobicity confirmed using web-based peptide analysis tool at www.peptide2.com/N_peptide_hydrophobicity_hydrophilicity.php) containing the Threonine residue (T993) required for the activation of LATS by HIPPO phosphorylation (T1079 in humans) (Supplementary Figure 2A) (Hergovich et al., 2006;Ni et al., 2015). The MOB binding motif is highly conserved in *Hvul*_LATS as compared to the human and mouse (Figure 2.4 B). The auto-activation T-loop (near S909 in human) of *Hvul*_LATS is 100 % conserved (Motif: AHSLVGTPNYIAPEVL) near S830 (Supplementary Figure 2B) (Ni et al., 2015). *Hvul*_MOB is highly conserved (84 % identity with human MOB) as compared to any other components of Hippo pathway homologs in *Hydra* indicating highly conserved function. The same site as reported for Human MOB is also highly conserved in *Hvul*_MOB at T35 (Motif: LLKHAEATLGSGNLR) (Supplementary Figure 2C). This site is crucial for the release of LATS-MOB complex from the MST-SAV-LATS-MOB complex and further initiation of LATS auto-activation (Ni et al., 2015). *Hvul*_YAP domain analysis revealed that it had a conserved TEAD-Binding Domain (TBD) and two WW domains. A serine phosphorylation prediction for YAP primary sequence was performed using GPS 2.1 web-based tool (Xue et al., 2010). Based on GPS prediction and manual curation, *Hvul*_YAP is predicted to have LATS phosphorylation site at S74 (motif: PIHTRAR**S**LPSNIGQ) and S276 (motif: YTAYMN**S**SVLGRGSS) homologous to the S127 (motif: PQHVRAH**S**SPASLQL) and S381 (motif: SDPFLN**S**GTYHSRDES) (Supplementary Figure 2D). Similar to mammals, a phosphodegron motif (DSGLDG) was identified immediately downstream to the S276 site (S381 in humans) which could be phosphorylated by CK1-ɣ at S287 (S388 in humans) of *Hvul*_YAP (Supplementary Figure 2E) (Zhao et al., 2010). These analyses indicated that the Hippo pathway effector protein YAP is well equipped for regulation by the LATS and CK1-ɣ. With its defined TEAD binding domain and WW domain, it could interact with transcription factor TEAD and other reported PPXY domain-containing proteins.

**Figure 2:**
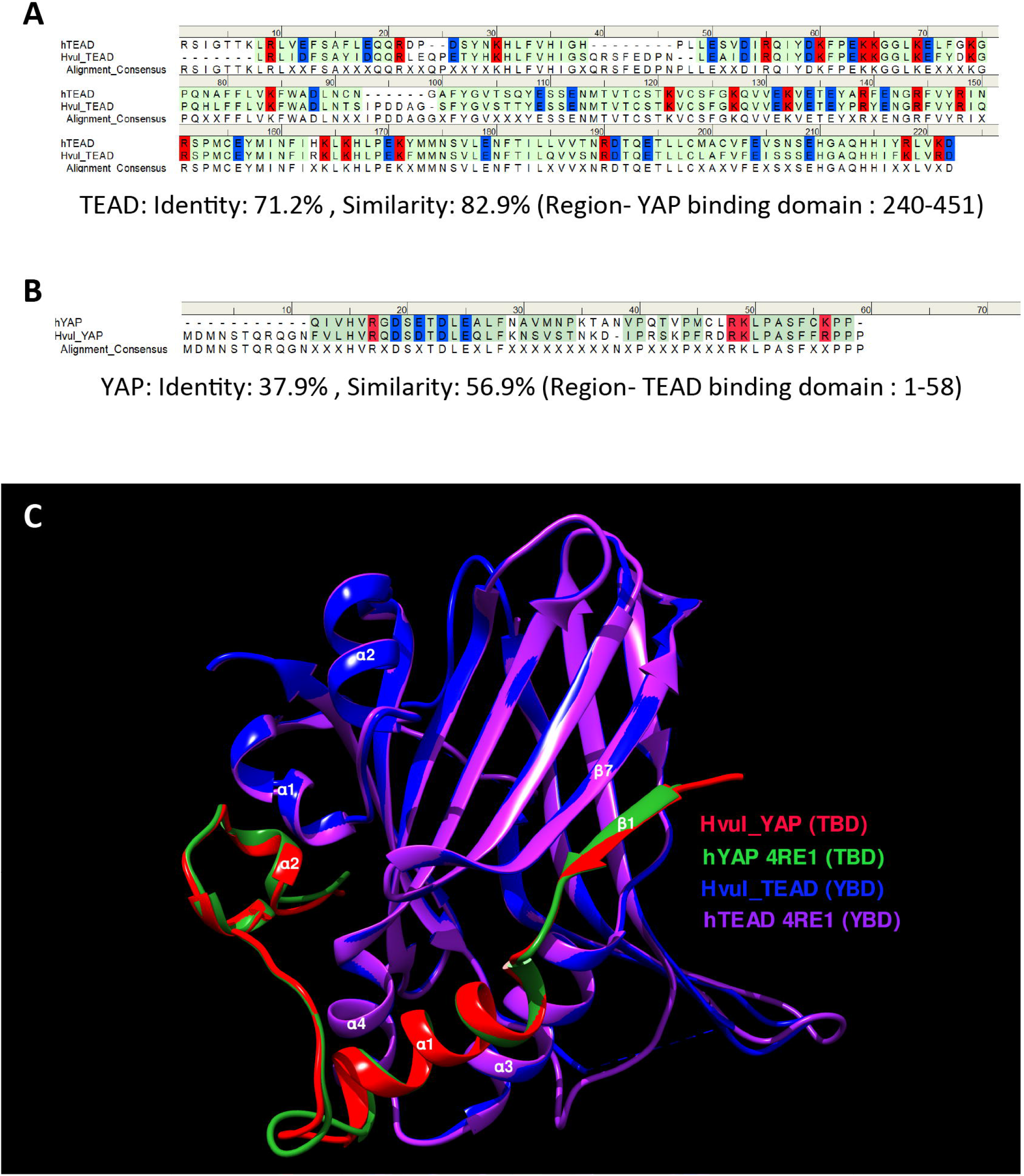
YAP-TEAD interaction domain is structurally conserved in *Hydra*. **A.** Sequence alignment of human and *Hydra* TEAD YAP-binding domains (YBD) showing 71.2 % sequence identity. **B.** Sequence alignment of human and *Hydra* YAP TEAD-binding domains (TBD) showing 37.9 % sequence identity. The alignment consensus shows conserved amino acid residues at a given position. If the there is no conservation, the position is labelled as X. Colour code: amino acid residues with positive charge-red, negative charge-blue and neutral-green. **C.** Structural superposition of predicted *Hvul*_TEAD YBD and *Hvul*_YAP TBD with YBD and TBD complex in Human (PDB:4RE1) showing highly conserved β-strands and α-helices structural placement. Important α-helices and β-strands are indicated with their number identification which are involved in the interaction of YBD and TBD. Colour code: Red- *Hvul*_YAP TBD, Blue- *HVUL*_TEAD YBD, Green-human YAP TBD (PDB ID-4RE1), Purple-human TEAD YBD (PDB ID-4RE1). D) Interaction of YBD (green color) with the TBD (purple color) in *Hydra* modelled using 4RE1 structure showing how the globular YBD (depicted in surface features) is bound by TBD (depicted as ribbon) through interactions at three different regions -region 1, region 2, and region 3. The amino acid side chains from TBD are represented as sticks for understanding their role in the interaction.

### Structural features of YAP and TEAD interaction

The Hippo effector protein YAP is known to elicit its biological function as transcription co-effector by interacting with transcription factors. Presently, YAP is known to interact with TEAD, β-catenin, SMAD, RUNX, p73 and ErbB4 for regulating their transcriptional responses as an activator or repressor (Strano et al., 2001; Komuro et al., 2003; Zhao et al., 2008; Szeto et al., 2016; Passaniti et al., 2017; Pan et al., 2018). Among these, YAP-TEAD interaction has been extensively studied and is known to be important for regulating cell growth and size as well as tissue architecture (Totaro et al., 2018). The interaction of YAP and TEAD was first shown to form through their specific interaction domains in 2001 (Vassilev et al., 2001). The structural features of this interaction in humans were first demonstrated in 2009 showing how the TEAD binding domain (TBD) in YAP (amino acids 53-99) interacted with the YAP binding domain (YBD) in the TEAD (position: amino acids 209-426) (Li et al., 2010). The YBD consists of 12 β strands which arrange themselves into two β sheets in an opposing fashion to form a β-sandwich fold. The four α helices from the YBD are arranged at the two ends of the β-sandwich fold for stabilizing the structure. The study showed that TBD-YBD interaction occurs over 3 interfaces. Each interface consisted of one of the following secondary structure of the TBD- the β1 strand, α1 helix or α 2 helix responsible for interacting with the globular YBD of the TEAD at the C-terminal. It was shown that the β1 strand of TBD interacted with the β7 strand of the YBD (interface 1), The α1 helix from TBD interacted with α3 and α4 helices of the YBD (interface 2). The α2 of the TBD was bound to the YBD through its interaction with α1 and α2 helices (interface 3) (Li et al., 2010).

Amino acid sequence alignment of the predicted YBD (amino acids 240-251) and predicted TBD (position: 1-58) of *Hvul*_YAP and *Hvul*_TEAD respectively with Human YAP and TEAD revealed 71.2 % sequence identity (82.9 % sequence similarity) of YBD (Figure 2 A) and a 37.9 % sequence identity (56.9 % sequence similarity) of TBD (Figure 2 B) which indicates plausible structural conservation and hence interacting capability of TBD with YBD. To confirm the same, the 3D structure of the YBD and TBD of *Hvul*_YAP and *Hvul*_TEAD was modelled using MODELLER software (Webb and Sali, 2016). The modelling was done based on the 4RE1 X-ray diffraction structure deposited at Research Collaboratory for Structural Bioinformatics PDB (RCSB PDB- https://www.rcsb.org/) which models the interaction of human homologs of TBD and YBD at a resolution of 2.20 Å. The model generated from the *Hydra* homologs was superimposed on the human YAP (hYAP) and hTEAD structure from 4RE1 and was found to highly structurally similar (RMSD for *Hvul*_YAP:hYAP- 0.338 A° and for *Hvul*_TEAD:hTEAD- 0.310 A°) and indicated a conserved interaction capability of *Hvul*_YAP and *Hvul*_TEAD (Figure 2 C). The modeled YBD-TBD complex of *Hydra* clearly shows how three different regions- Region 1, Region 2 and Region 3 of TBD (purple) interacts with the globular YBD (green) by non-covalent bond interactions (Supplementary Figure 3A). The Region 1 interface consisting of TBD β1 (amino acids 10-17) and YBD β7 (358-363) strands interact with seven hydrogen bonds in the human complex, forming an anti-parallel β sheet (Li et al., 2010). In *Hydra* there are only six hydrogen bonds (green dotted lines) due to the presence of Gln18 in β1 instead of Gly59 found in humans (Li et al., 2010), introducing a rotation in the preceding Arg which disables it from forming a hydrogen bond (Supplementary Figure 3B). The 2^nd^ interface (Region 2) has the α1 helix of the TBD (amino acids 20-32) fitting right into the binding groove of the YBD formed by the α3 and α4 helices of the YBD (amino acids 385-409) (Supplementary Figure 3C). Similar to humans; this region is mainly mediated by hydrophobic interactions with the α1 helix of the TBD having conserved LXXLF motif for hydrophobic groove binding (Li et al., 2010). This interaction mainly consists of Leu24, Leu27and Phe28 from TBD and Try386, Lys393 and Val406 of YBD (pink dotted lines). In *Hydra*, few hydrogen bonds (green dotted lines) not found in humans may lead to a more stable interface. The 3^rd^ region (3^rd^ interface) consists of a twisted coil and α2 helix (amino acids 42-58) from the TBD interacting deeply with the pocket formed by the α1 helix, β4, β11 and β12 helices of the YBD. This region was found to be indispensable for the YAP-TEAD complex formation in humans (Li et al., 2010). The region 3 in *Hydra* contains the hydrophobic side chains of the TBD – Phe44 (Met86 in humans), Leu49, Pro50 and Phe53 forming extensive van der Waals interactions with the YBD of TEAD at Glu280, Ala281, Ile282, Gln286, Ile287, Leu312, Leu316, Val431, His444 & Phe446 (Supplementary Figure 3D). The interface is further strengthened by multiple hydrogen bonds (indicated in green dotted lines) – TBD_Arg47:YBD_Gln286, TBD_Lys48:YBD_Gln286, TBD_Ser52:YBD_Glu280 and YBD_Lys314:TBD_Phe53. The hydrophobic interactions in region 3 consists of Phe53, Pro50 and Phe44 from TBD and Lys314, Glu408 and Phe446 from the YBD. While the hydrophobic interactions involving Pro56 and Pro57 from TBD with Trp316 and His444 respectively help to push the proline residues out of the hydrophobic pocket. One of the unique aspects that can be predicted from the model is that *Hydra* region 3 YAP-TEAD complex is able to form two salt bridges (orange dotted lines) - TBD_Arg47:YBD_Asp289:YBD_Asp289 and TBD_Lys48:YBD_Asp283:YBD_Asp451. The human complex only forms a salt bridge at TBD_Arg89:YBD_Asp249: YBD_Asp249. These observations indicate a more stable YAP-TEAD interaction in *Hydra* as compared to the humans. To shed more light on the same, the computationally calculated binding energy of the YAP-TEAD complex between the two organisms were compared using the web-based server PRODIGY (PROtein binDIng enerGY prediction) in Protein-protein mode (Xue et al., 2016). The ΔG of YAP-TEAD complex in humans is about −6.8 kcal mol^−1^ while the complex in *Hydra* has a value of −14.7 kcal mol^−1^. This large difference in the binding energy supports the possibility that the YAP-TEAD complex in *Hydra* is much more stable.

**Figure 3:**
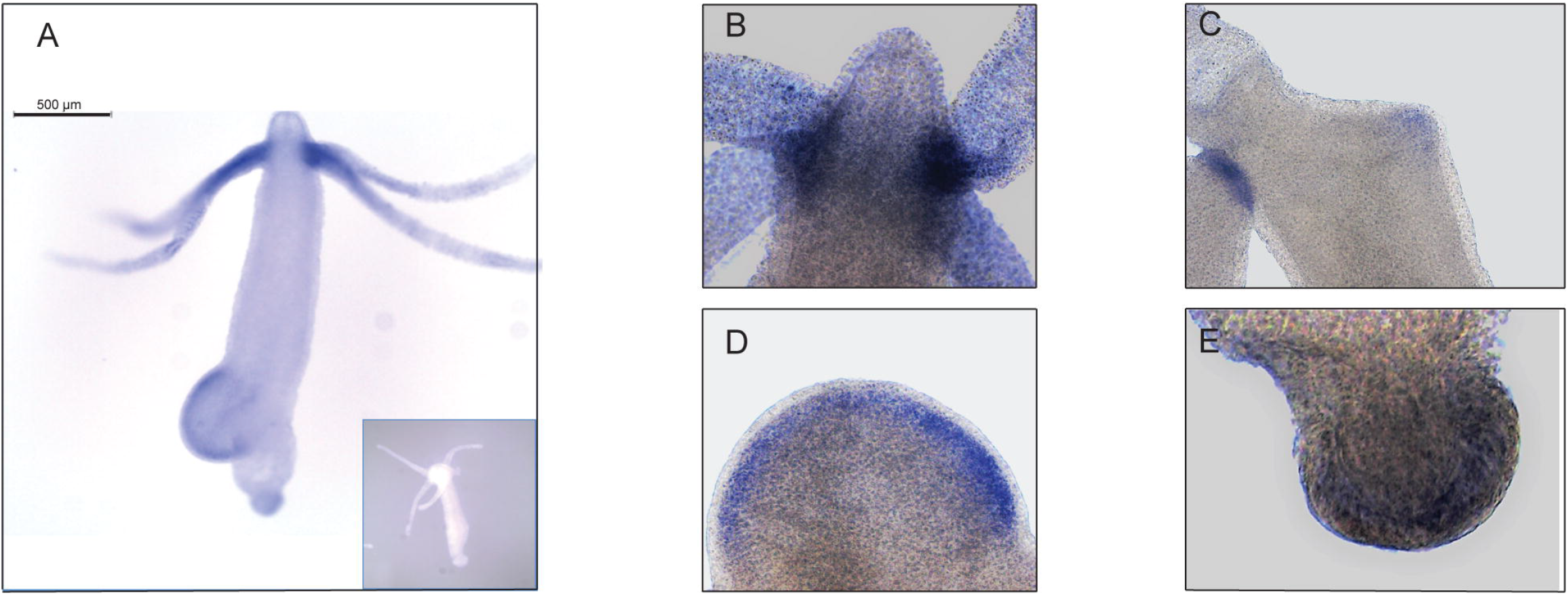
*Hvul_yap* expression analysis in *Hydra*. Whole-mount in situ hybridization of *Hvul_yap* expression at **A.** Region across the polyp (inset shows polyp probed with a sense RNA probe). **B.** Head, **C.** basal disk, **D.** mid-stage bud, **E**. early bud/ late bud foot. The scale bar is 500 μm long.

### Expression analysis of the Hippo pathway genes in *Hydra*

#### *Hvul_yap* expression in *Hydra*

The expression pattern of *Hvul_yap* in *Hydra* polyp was studied by whole-mount *in situ* hybridization (WISH). The staining pattern observed from the whole polyp indicates low-level expression throughout the body with higher expression at the tentacle base and tip of the early stages of the developing new bud (Figure 3A). A closer look indicates that the expression is stronger in the endodermal cells as compared to the ectodermal cells (Figure 3B - E). *yap* expression in the early stages of bud development indicates its role in budding. Higher *yap* expression at the region of high mechanical stress such as the tentacle base, early budding tip and mature bud-parent polyp boundary indicates a probable ancient mechano-sensory role of YAP in *Hydra*. These polyps were cryosectioned to obtain a closer look at the types of cells expressing *yap* (Supplementary Figure 4). The images of these sections revealed cells in doublets, quadruplets and groups of cells among other stained cells indicating their interstitial stem cell origin, plausibly nematoblast and nests of nematoblasts.

### Expression pattern of *Hvul_hpo, Hvul_mob* and *Hvul_sav* genes

An RNA WISH study of *Hvul_hpo* showed expression throughout the gastric region (Figure 4A). No expression was observed at the differentiated zones of hypostome, tentacle or basal disk which might indicate a role in stem-cell maintenance or differentiation but not in terminally differentiated cells. There is a slight reduction in expression at the budding zone and early buds which might indicate the antagonistic role of HPO towards YAP activity in areas of high mechanical stress as reported in other organisms. *Hvul_hpo* expression can also be seen at mature bud-parent polyp boundary indicating a fine-tuning of regulation of Hippo pathway-dependent during bud detachment. *Hvul_mob* expression showed a similar pattern to that of *Hvul_yap* with a distinct down-regulation at the basal disk region of both adult and budding *Hydra* (Figure 4B). *Hvul_sav* expression reflected the expression pattern of *Hvul_hpo* indicating a similar role. It can also be noted that there is marked reduction in expression at the budding region, early and late buds, unlike the *Hvul_hpo* (Figure 4C).

**Figure 4:**
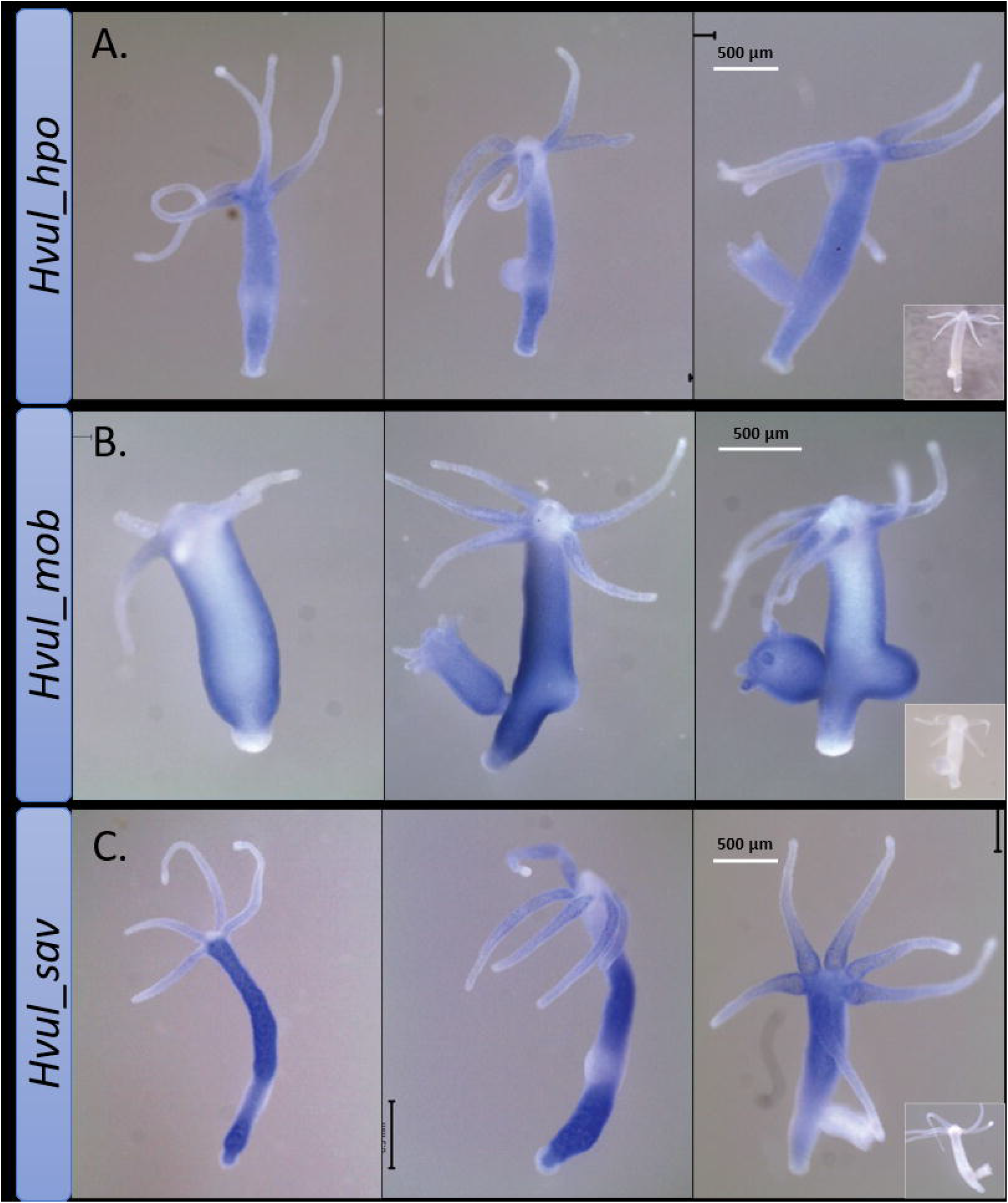
*Hvul_hpo*, *Hvul_mob* and *Hvul_sav* expression analysis in *Hydra*. Whole mount in situ hybridization of **A.***Hvul_hpo*, **B.***Hvul_mob* and **C.***Hvul_sav* expression for the whole polyp. The insets on the right indicate negative controls probed with sense RNA probe. The scale bar is 500 μm long.

A recent study reported high-throughput sequencing of the transcriptome of 24,985 single *Hydra* cells using Drop-seq and identified the molecular signatures of various cell states and types (Siebert et al., 2019). The differential expression of the Hippo pathway components and their pattern were examined using the Single Cell Portal. The expression patterns of *Hvul_yap, Hvul_tead, Hvul_hpo, Hvul_lats, Hvul_sav & Hvul_mob* were queried. From the single-cell data, *Hvul_yap* expression was found to be insignificantly dysregulated or differentially expressed between cell-types (Supplementary Figure 5). Surprisingly, such a trend was commonly observed between all the other Hippo pathway components namely, *Hvul_tead, Hvul_hpo, Hvul_lats, Hvul_sav & Hvul_mob*. This indicates a slight disparity with the WISH data. This could be due to lack of enough resolution from the datasets used. The data showed here only represent a relative fold-change between the cells and may indicate that the expression levels are relatively the same between the cells. The WISH data also indicate that most of the cells express almost all types of Hippo pathway components, yet at the same time, we see that they are excluded from some regions. These may be extremely stage-specific and hence difficult to be picked up in sc-RNAseq of whole polyps. Nevertheless, the findings from analyzing single-cell data argue in favor of the fact that the Hippo pathway components are essential for cells and need to be expressed in almost all cell-types. Their activity might be regulated at the protein level, and hence a protein-based analysis is essential to better understand the regulation of Hippo pathway in *Hydra*.

### Protein Expression analysis of the *Hvul*_YAP in *Hydra*

#### Region-specific and cell-type expression of *Hvul*_YAP in *Hydra*

The *Clytia hemispherica* specific Yorkie (CheYki) antibody was raised against a peptide from the WW1 region of CheYorkie in rabbit (Coste et al., 2016). The CheYki peptide sequence was extracted from the Marine Invertebrate Model Database (MARIMBA) and was used to align with *Hvul*_YAP using CLUSTAL Omega. The full protein alignment showed just a 39.36 % identity. However, a peptide-specific (immunogen) alignment gave a 60 % identity which raised the probability of cross reactivity of this antibody against *Hvul*_YAP (Supplementary Figure 6 A). To test the same, an immunofluorescence assay (IFA) was run using CheYki antibody or pre-immune serum. The IFA yielded a robust signal for CheYki antibody as compared to the negative control (Supplementary Figure 6 B). Examination of localization of YAP expressing cells revealed a pattern similar to what we found in YAP ISH (Figure 5 A). The expression was seen more or less throughout the body. The base of the tentacle showed high expression similar to that seen in ISH but the number of YAP expressing cells drops in hypostomal region and the inter-tentacle zone. Unlike the pattern of transcripts seen in the ISH, the YAP expressing cells were depleted at the basal disk region.

**Figure 5:**
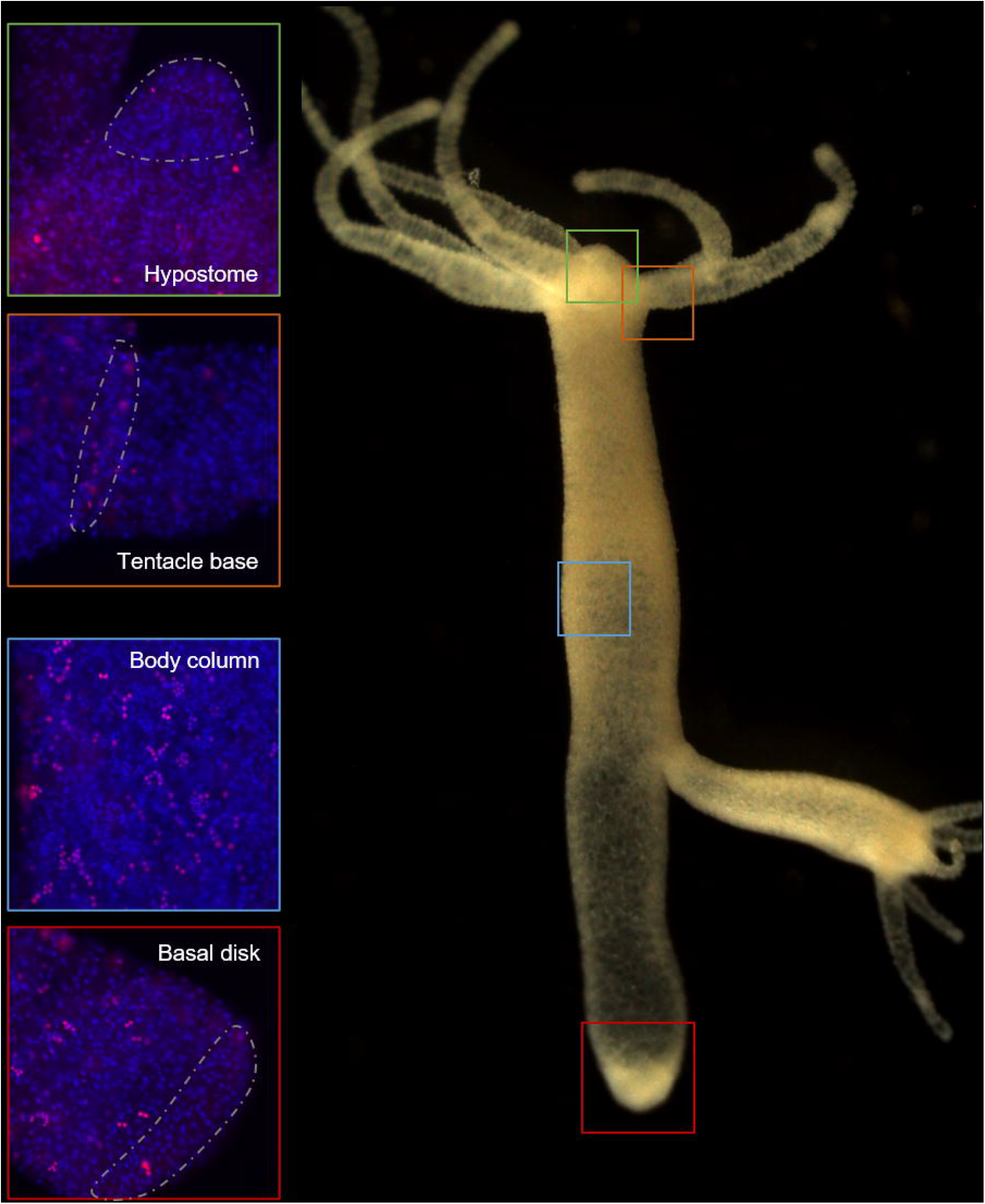
Expression of *Hvul*_YAP in *Hydra*. Immunofluorescence assay of *Hvul*_YAP performed using anti-CheYki antibody showing localization of YAP positive cells at various locations in an adult polyp. The red fluorescent dye shows Alexa 594 staining of YAP and the blue dye shows DAPI staining of nucleus. The hypostomal region is indicated by a green box, the tentacle base is indicated by an orange box. The body column is indicated by a blue box and basal disc area is indicated by a red box.

The body column of *Hydra* is uniformly interspersed with YAP expressing cells (Figure 6). These cells can be seen almost exclusively in groups (duplets, quadruplets or more). There were specific patterns of these groups which looked similar to the ones seen in cryosections of ISH samples (Supplementary Figure 4). The expression was clear for nuclearized YAP while the cytoplasmically localized YAP were dispersed and difficult to observe. A careful analysis of cells expressing YAP based on the staining intensity and intercellular distance, as seen in IFA indicates different subsets of cells. Based on the YAP expression intensity, there seem to be cells exhibiting high expression (Blue arrow), medium expression (yellow arrow) and low expression (green arrow). Based on the cellular clustering, cell types can be divided into cells which are duplets or quadruplets (orange arrows) which may be interstitial stem cell undergoing first and second mitotic division. There are also clusters of cells which are arranged into a linear file whose identity is difficult to judge (red arrows). Yellow arrows indicate clusters of cells which looks like part of a nest of nematoblasts. These nest cells are typically arranged into 8-16 cell-clusters. As can be noticed here, these clusters are not completely YAP expressing, and only a subset of these express YAP. This may indicate that these cells are expressing only at certain stages of nematoblast differentiation. Such similar clusters can be observed even in the high-level YAP expressing cells (blue arrows) indicating a yet different subset of nematoblast cells. A different population of cells shows extra-nuclear staining (white arrow). These stains might be non-specific since they are localized in cysts similar to that seen in desmonemes and stenoteles. These results suggest that at least some of the YAP expressing cells have interstitial cell origin. While inner hypostome (area immediate around the mouth) and the tentacles are virtually devoid of YAP expressing cells, we find that there are a few non-clustered YAP expressing cells at the region interstitial to the tentacle bases and the outer hypostome (Figure 5 and Supplementary Figure 7B). Such an expression pattern may indicate a role of YAP in tissue compartment-boundary regulation for hypostomal and tentacle development and/or maintenance of gene networks in *Hydra*. Such a role of Yki (YAP) has been recently proposed in *Drosophila* in wing imaginal disc development by regulating the expression of Hox genes and Hedgehog signaling (Bairzin et al., 2020).

**Figure 6:**
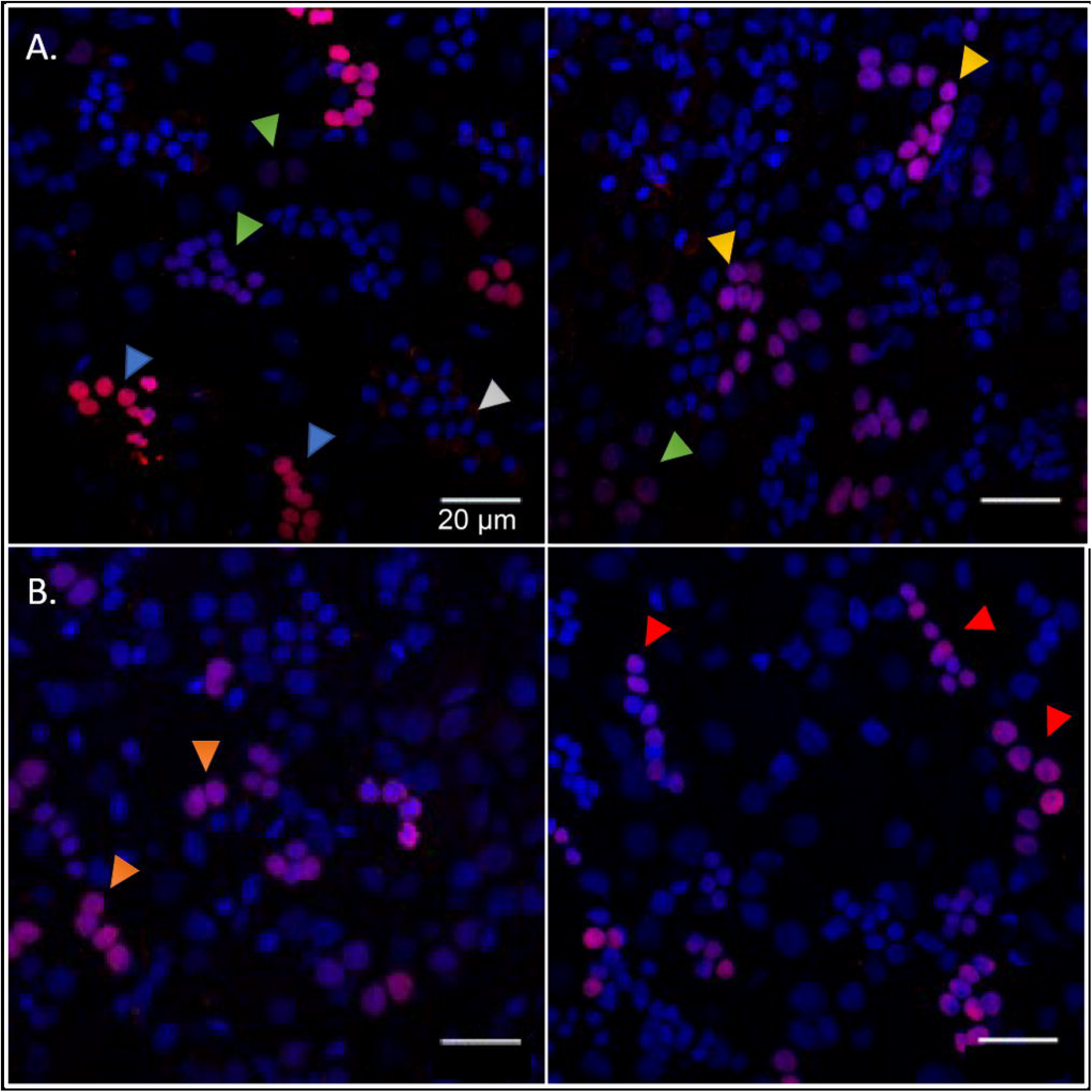
Types of *Hvul* _YAP expressing cells in *Hydra*. Immunofluorescence assay of *Hvul*_YAP performed using anti-CheYki antibody on macerated cells at 60X. **A**. This panel shows cell types based on signal intensity or YAP expression level in cells. Blue arrow represents cells with high YAP expression, yellow arrow represents cells with medium YAP expression, cells with a green arrow represents low YAP expression. White arrow indicates extra-nuclear staining in nematocysts. **B**. This panel depicts cell types based on the cellular arrangement. Orange arrows represent cells with duplet or quadruplet arrangement and red arrow represents cells arranged linearly. Red: YAP & Blue: Nucleus (Magenta indicates merged image). Immunofluorescence assay using the anti- *Hvul*_YAP antibody of macerated cells at 60X. The red fluorescent dye shows Alexa 594 staining of YAP and the blue dye shows DAPI staining of nucleus. (Scale bar: 20 μm)

### YAP expressing cells are recruited to newly developing buds but are excluded from the hypostomal region upon initiation of differentiation

Cellular dynamics of YAP expressing cells during *Hydra* bud development was studied using immunofluorescence assay. Buds at different points of bud-development from early to late stages were observed (Stage 3, 4, 6 and 9). It was clear that the YAP expressing cells moved into the early bud with an expression pattern very similar to that found in the body column. Such pattern is persistent throughout the budding stages in the body column of the newly developed bud. The most interesting changes happening to the YAP expressing cells in a bud is at the hypostomal region. The YAP expressing cells near the distal bud tip were found to be non-clustered as compared to the rest of the lower bud region. At stage 3, the bud-tip where the head organizer has been set for establishing the new body axis for bud, YAP expressing cells seems to be depleted (Figure 7A). This pattern is even more conspicuous from stage 4 onwards (Figure 7B-D, Supplementary Figure 7A). From stage 9 onwards, the expression pattern similar to the adult *Hydra* is established where we see non-clustered YAP expressing cells seen sparsely at the boundaries between the hypostome and the tentacle base (Figure 7D, Supplementary Figure 7B). The appearance of non-clustered cells in these regions may indicate a different sub-type of YAP expressing cells having a role in head organizer maintenance in *Hydra*. Another interesting point to note is that YAP expressing cells are completely depleted at the basal disk (Figure 5), hypostome and tentacles. This observation may indicate an important antagonistic role of YAP signaling in tissues with terminally differentiated cells. The lack of YAP expressing cells even at the early developmental stages of tentacle development in a new bud (Supplementary Figure 7B) and at the Adult-bud boundary where the basal disk will form (Supplementary Figure 7C) further suggests the possibility of Hippo pathway in cell differentiation.

**Figure 7:**
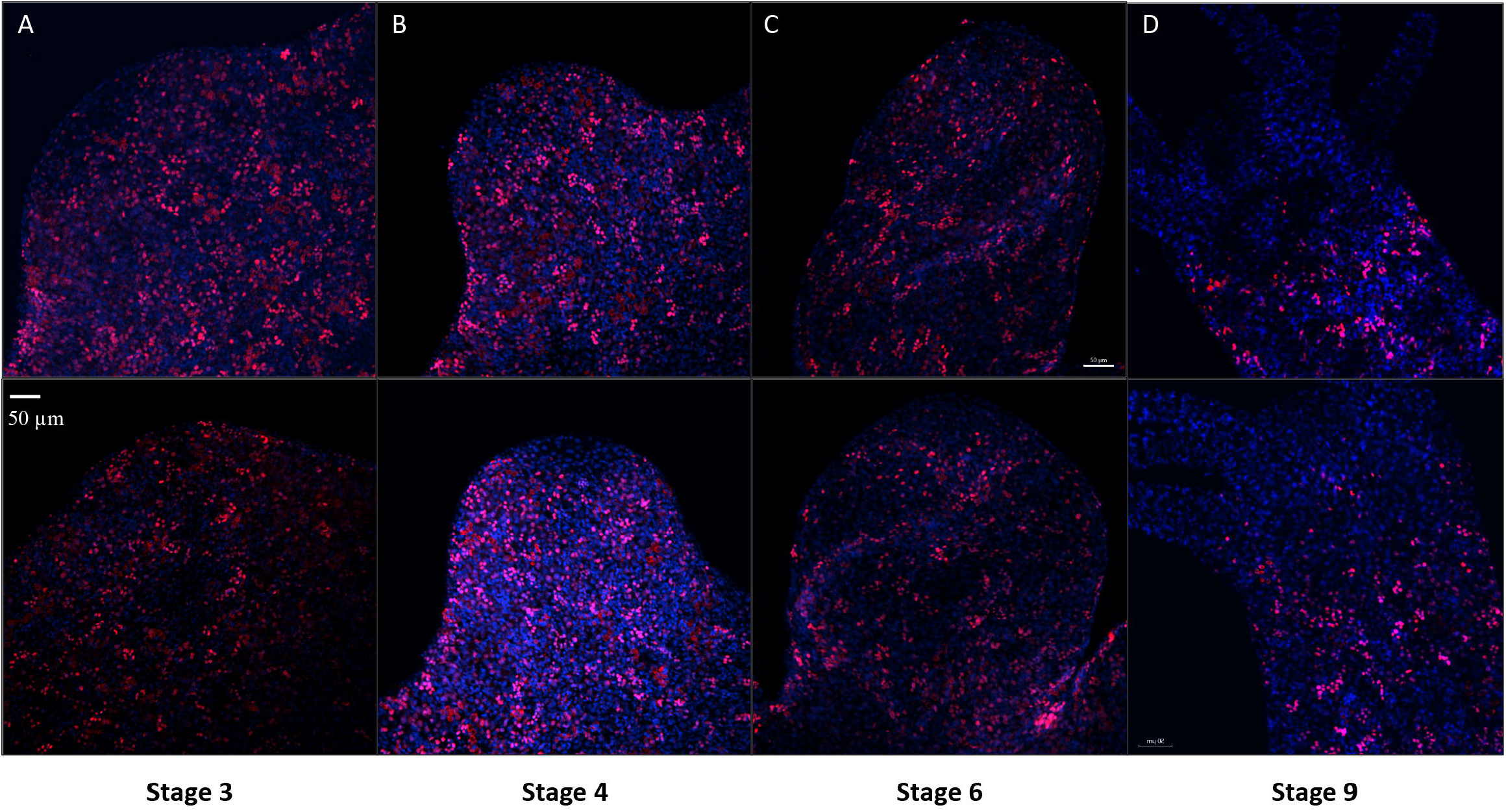
YAP positive cells are recruited early to the bud tip and are excluded from the region which are terminally differentiated in the late stages of bud development. Immunofluorescence assay of *Hvul*_YAP performed using anti-CheYki antibody for different budding stages (represented by two polyps for depicting each stage) of *Hydra* showing recruitment of YAP positive cells to the budding tip. **A.** Stage 3 shows early recruitment of YAP positive cells to the emerging bud with non-clustered cells at the distal tip with slight depletion at tip of the bud. **B.** at Stage 4, depletion of the YAP expressing is more prominent which gets further exaggerated at **C.** Stage 6 and **D.** Stage 9. Red: YAP & Blue: DAPI. (Scale bar = 50 μm)

### YAP expressing cells are early responders to head amputation

Immunostaining for YAP on decapitated polyps shed light on the participation of YAP expressing cells during early regeneration (Figure 7). YAP expressing cells can be observed occupying the site of injury within one hour of amputation. These cells increase in density as time progresses until 4 hours post-amputation (hpa). Since wound healing takes approximately 1-2 hpa, the YAP expressing cells may migrate along with the epithelial cells during the wound-healing phase. An increase in density might be caused by either further migration of the YAP expressing cells from the body column or by division from the pre-existing cells at the site of injury. The interesting observation to note is that the population of YAP expressing cells at the site of injury until 4 hpa is similar to the population seen in the body column of *Hydra* (green arrows). This changes after 8 hpa as the YAP expressing cells at the distal-most region of the regenerating tip assumes a new cell-type characteristic. These cells are non-clustered and look similar to the cells observed at the budding tip and at the region around the boundaries of hypostome and tentacle base in adult *Hydra*. They can now be seen as individual cells arranged arbitrarily at the tip. Since YAP is a known mechanotransducer, these cells are either differentiated from the cells migrated from the body column or are cells assuming a new phenotype in response to the mechanical change in the cellular environment due to lack of ECM and physical disruption of cells (Shimizu et al., 2002). The dynamics of YAP expressing cells in the regenerating tips indicate that they are recruited to the site of injury early during the regeneration and are probably early responders of mechanical changes.

## Discussion

Detailed characterization of the Hippo pathway and its components in pre-bilaterians has been extremely sparse. There have been few studies reporting the presence of Hippo homologs in these primitive organisms. *Capsaspora owczarzaki*, a single-celled eukaryote is the most primitive organism predicted to have a complete set of functional core Hippo pathway homologs indicative of a holozoan origin of the functional pathway (Sebé-Pedrós et al., 2012). Another study confirmed the presence of Hippo pathway components in a Ctenophore species: *Pleurobrachia pileus* and a Cnidarian species *Clytia hemispherica* (Coste et al., 2016). While this study reported an absence of Yki in Ctenophores, it showed that the Yki in *Clytia* are conserved for the regulation of cell proliferation and growth. In this study, we have for the first time identified and characterized a complete set of core Hippo pathway components in *Hydra vulgaris* through bioinformatic analysis and cloning. The current phylogenetic analysis is in congruence with previous report that *Nematostella vectensis* homologue is more similar to complex vertebrates (Hilman and Gat, 2011). In fact, all the Cnidarian homologues exhibit higher similarity with the chordate YAP sequences. This suggests that YAP sequences evolved close to the emergence of the chordate homologs and might exhibit similar properties observed in these organisms. Domain organization analysis indicates divergence in the N-terminal homology domain of YAP (FAM181) in different lineages. This suggests the clade specific role of the FAM181 region, probably in the interactions with TEAD like or other proteins. This further indicates taxon-specific modification took place in the FAM181 region and might play lineage specific functions. We show that the Hippo pathway components are more or less uniformly expressed throughout the polyp tissues barring a few regions in a gene-specific manner like budding zone, early buds, extremities of the polyps such as tentacle tips or basal disks. Considering the studies in bilaterians indicating that these components are all tightly controlled to regulate the cell cycle and cell differentiation, it can be easily seen why these genes are expressed uniformly in all tissues. Since the extremities of the polyps are terminally differentiated, they probably do not need these genes for the functions mentioned above and are already set to perform its designated functions without needing any change. The analysis of amino acid sequences of these genes to predict the secondary structure and 3D tertiary protein models have also given us some insightful results. Domain architecture of all the Hippo pathway proteins shows that their architecture is well conserved in Cnidaria, which confirms an ancient establishment and evolution of the pathway in the basal metazoans. The 3D modelling of YAP’s TBD and TEAD’s YBD in *Hydra* using the published crystal structure of their Human homolog predicts a similar interaction capability of YAP and TEAD in *Hydra*. Our analysis revealed that the YAP-TEAD complex is highly stable in *Hydra*. This raises the possibility that the YAP-TEAD interaction was robust in primitive metazoans, and as the signaling pathway evolved, the stability of the complex was presumably partially compromised to accommodate the promiscuous nature of YAP in more complex organisms. This indirectly indicates that the functions of the Hippo pathway or YAP signaling reported in bilaterians may have been established as early as in Cnidarians and hence may have played in developing important characteristics of multicellular organisms like cell-type divergence, body-axis development, germ-layer differentiation etc.

In *Clytia*, it was found that Yki was nuclearized at the tentacle base where there are highly proliferating cells, while they are inhibited in the tentacles where the cells are differentiated (Coste et al., 2016). Using the antibodies used in the same study, we were able to study the protein-level expression of YAP in *Hydra*. We find that even though *Hvul_yap* is expressed uniformly throughout the polyp, only a few cells have *Hvul*_YAP in the “active form” (nuclearized). We find that these nuclearized YAP are more or less uniformly spread throughout the polyp. YAP expression is almost absent or not nuclearized in the terminally differentiated regions including the tentacles, hypostome or basal disk. An interesting observation is the presence of YAP expressing cells at the tentacle base forming a circle (Figure 5 and Supplementary Figure 7B). This can be considered homologous to the expression pattern seen in *Clytia* which may be speculated as necessary for terminal differentiation of cells while crossing the body column-tentacle boundary. Another possibility can be the mechanical activation due to physical stress experienced at the tentacle base due to movement of tentacles or anatomical constraints. Most of these cells in the body column can be seen in groups or colonies. Cellular features and arrangements of YAP positive cells are indicative of interstitial stem cell origin. Cells like desmonemes and stenoteles are mechano-sensitive, and YAP may regulate their development and function. Another interesting observation is the presence of a non-clustered group of cells in the outer hypostomal region (Supplementary Figure 7B). Such an expression pattern raises many interesting possibilities. It is reported that the ectodermal cells in the hypostome is maintained separately from the gastric region (Dübel et al., 1987;Dübel, 1989). The inner hypostomal ring consists exclusively of terminally differentiated cells (Dübel, 1989). The stationary region in the hypostome (the outer hypostomal ring) contains a population of the ectodermal epithelial cells that retains its proliferative potential which contribute exclusively to the cell-types in the entire hypostomal region. Once the hypostome is specified, there are no contributions from the gastric ectoderm towards hypostomal cells unless the hypostome is lost upon amputation. A unique population of YAP expressing cells (non-clustered cells) in the outer hypostomal region and not at the inner region may indicate the possibility of these cells being maintained in their undifferentiated proliferative stem cell state by YAP for specific hypostomal functions. An identical population of cells can be found at the regenerating tip (Figure 8-white arrows) and during early bud development. These observations raise the possibility of these cells having a crucial role in establishing and maintaining the head organizer. This regulation may well be mechanically activated upon amputation or biochemically with pre-existing cues. The appearance of the non-clustered cells at the regenerating tip at 8 hrs is interesting since the expression of one of the variants of *brachyury* (Hy*Bra2*) coincides with the same time point in the regenerating *Hydra* ((Bielen et al., 2007); Unni et al., unpublished findings).The same study also shows early expression of *Bra* during bud formation. Interestingly, the expression pattern of *HyBra* is exclusively at the hypostomal region encompassing both the outer and inner hypostome (Technau and Bode, 1999; Bielen et al., 2007). *Bra* is known to be a direct responder to consolidation Wnt/β-catenin signaling (Yamaguchi et al., 1999). *HyBra* has also been implicated in the establishment of the head organizer in *Hydra* (Technau and Bode, 1999). Hence, this may mean that appearance of *HyBra* may coincide with the true setting up of the head organizer. This raises the enticing prospect of YAP expressing cells at the outer hypostome region to restrict the head organizer-related function of Brachyury to the inner hypostome ring by exerting its tissue boundary regulation functions via the hedgehog pathway (Bairzin et al., 2020).Taken together with the expression pattern of YAP in developing bud, adult polyp and the regenerating tip, consolidates the possibility of YAP in the establishment and maintenance of the head organizer function in *Hydra*.

**Figure 8:**
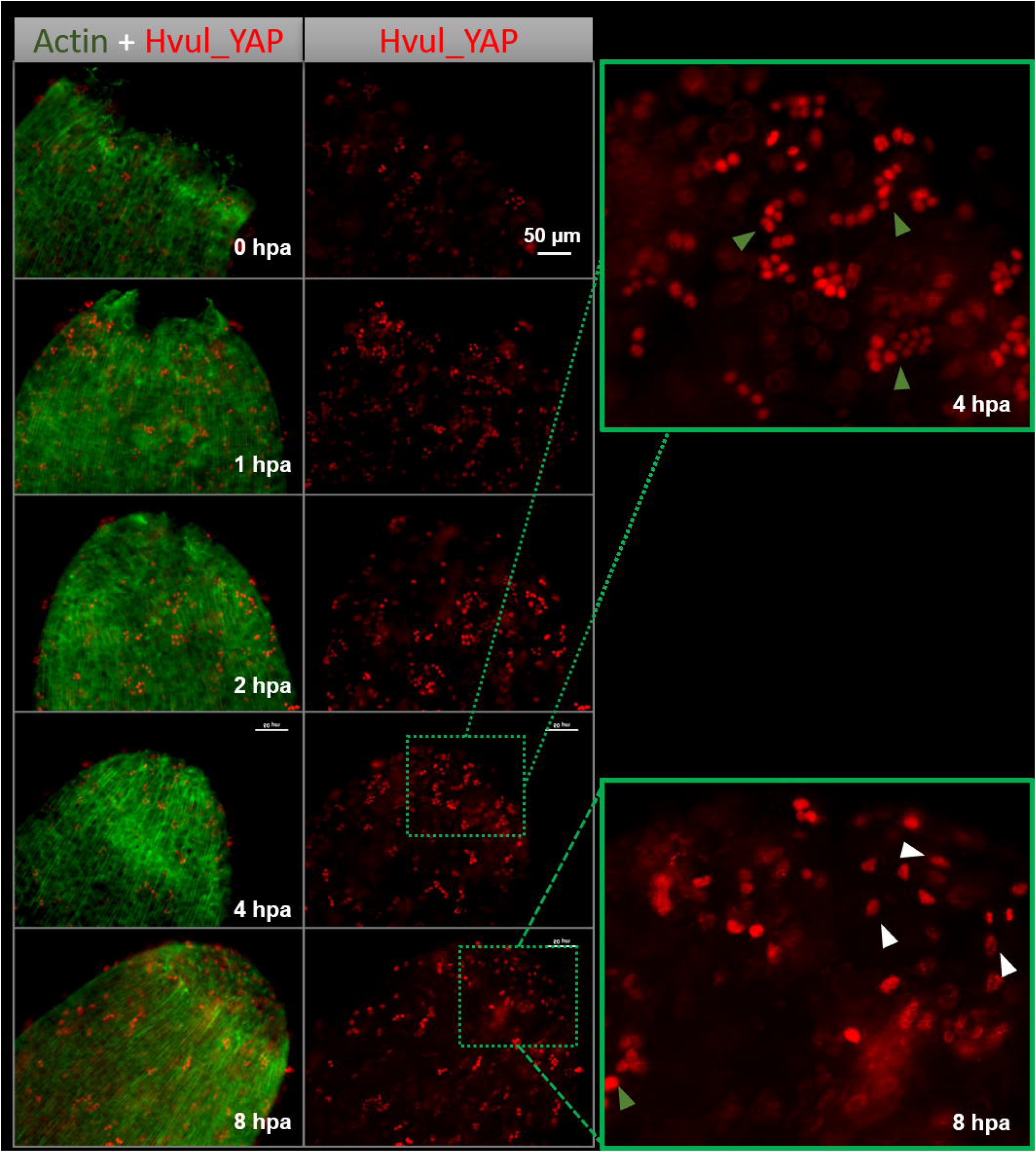
YAP positive cells are recruited early to the regenerating tip in *Hydra*. Immunofluorescence assay of *Hvul*_YAP performed anti-CheYki antibody for head regenerating *Hydra* showing recruitment of YAP positive cells to the regenerating tip. The density of YAP positive cells can be seen increasing at the site of injury from 1 hr post-amputation (1 hpa). White arrows indicated in the zoomed-in image shows the generation of a new type of YAP expressing cells at the regenerating tip by 8 hpa as compared to cell population seen away from the tip or previous time points (green arrows) Red: YAP & Green: Actin. (Scale bar = 50 μm)

This study shows that the Hippo pathway is an important signaling pathway capable of regulating the cellular differentiation and tissue regeneration in *Hydra*. A more in-depth study of YAP signaling under these contexts might reveal interesting insights into the evolution of the functions associated with complex organisms. YAP can act as a mechanotransducer and has been shown to play a role in regulating various morphogenetic and developmental functions. This aspect of YAP is only starting to be fully understood and have been poorly studied in basal metazoans to understand its origins. A detailed study in *Hydra* to understand the same will shed light on the fundamental aspects of how tissue mechanics plays a role in regulating cell function.

## Conflict of Interest

The authors declare that the research was conducted in the absence of any commercial or financial relationships that could be construed as a potential conflict of interest.

## Author Contributions

Conceptualization: M.K.U., P.C.R., S.G.; Methodology: M.K.U., P.C.R., S.G; Validation: M.K.U., P.C.R., S.G; Formal analysis: M.K.U., P.C.R., S.G.; Investigation: M.K.U. & P.C.R.;

Resources: S.G.; Writing - original draft: M.K.U., P.C.R., S.G.; Writing - review & editing: M.K.U., P.C.R., S.G.; Visualization: M.K.U., P.C.R., S.G.; Supervision: S.G.; Project administration: S.G.;

Funding acquisition: S.G.

## Funding

This work was supported by the Centre of Excellence in Epigenetics program (BT/01/COE/09/07) of the Department of Biotechnology, Government of India and the JC Bose National Fellowship from the Science and Engineering Research Board (JCB/2019/000013) (S.G.). The authors acknowledge funding from IISER Pune - intramural (S.G); Department of Biotechnology postdoctoral fellowship (P.C.R); and Early Career Fellowship (IA/E/16/1/503057) (P.C.R.); fellowships from the University Grants Commission (UGC) (M.U.); EMBO Short-term fellowship and Infosys Foundation for international travel support (M.U.)

## Acknowledgments

Authors wish to thank Dr Michaël Manuel for providing the kind gift of CheYki antibody, and Dr Inna Solomonov for useful comments on the manuscript and Dr Neeladri Sen for his help with protein modelling. We thank Prof Irit Sagi to allow use of facilities in her lab to perform *Hydra* YAP budding immunofluorescence assay and Mr Assaf Hanuna for maintaining *Hydra* culture at the Weizmann Institute of Science. We would like to thank Dr Yoseph Addadi for assistance with imaging samples using Andor Dragonfly Spinning Disc Microscope, Dr Rachel Paul for assistance with whole-mount RNA *in situ* hybridization of *Hvul_mob*, and the IISER-Pune Microscopy facility.

## Data Availability Statement

The datasets for this study can be found in the Supplementary Table 1. This includes the protein sequences used for phylogenetic analysis, the mRNA sequences and protein sequences of the *Hydra* Hippo pathway core homologs in separate tabs of the excel sheet.

## Notes

### Competing Interest Statement

The authors have declared no competing interest.

