## Supplementary material for "The Hippo pathway transcriptional co-activator YAP is involved in head regeneration and bud development in *Hydra*"

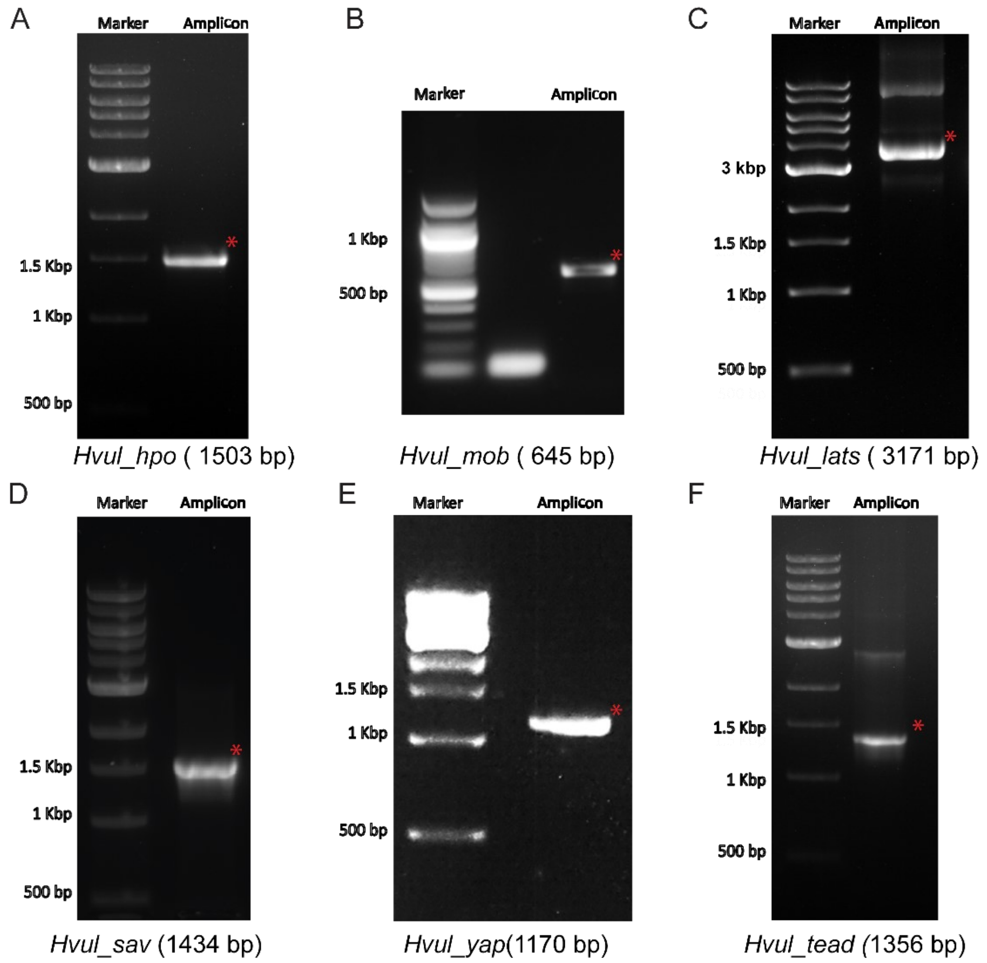


**Supplementary Figure 1. Validation of the Hippo Pathway homologs in *Hydra* by PCR mediated amplification using the predicted sequence**. **(A)** *Hvul*_Hpo was amplified at 1503 bp. **(B)** *Hvul*_Mob was amplified at 645 bp. **(C)** *Hvul*_LATS was amplified at 1170 bp. **(D)** *Hvul*_SAV was amplified at 1434 bp. **(E)** *Hvul*_YAP was amplified at 1170 bp. **(F)** *Hvul*_TEAD was amplified at 1356 bp.

A

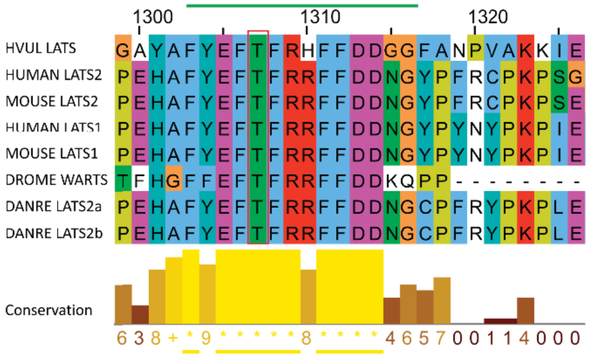


B

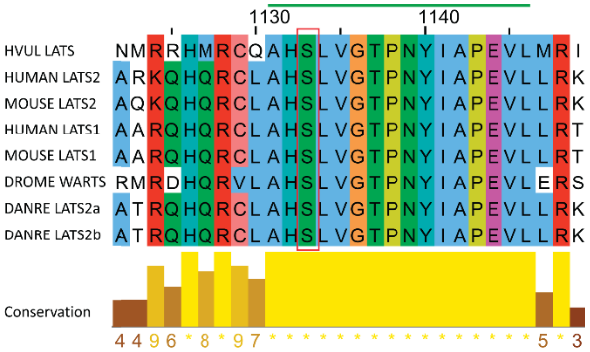


C

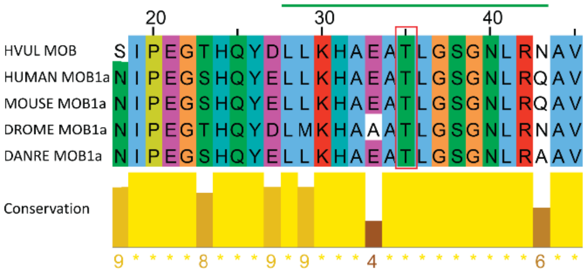


D

| ID | Position | Code | Kinase | Peptide | Score | Cutoff | Source | Logo |
|---|---|---|---|---|---|---|---|---|
| HVUL_YAP | 74 | S | AGC/Akt | PIHTRARSLPSNIGQ | 12.03 | 8.272 | Pred. | |
| HVUL_YAP | 277 | S | AGC/DMPK/CRIK | YTAYMNSSVLGRGSS | 0.545 | 0.463 | Pred. | |

E

| ID | Position | Code | Kinase | Peptide | Score | Cutoff | Source | Logo |
|---|---|---|---|---|---|---|---|---|
| HVUL_YAP | 287 | S | CK1/CK1/CK1-A/CSNK1A1 | GRGSSVDSGLDGMES | 24 | 21.704 | Pred. | |

**Supplementary Figure 2. Motif analysis of LATS, MOB and YAP. (A)** Conserved hydrophobic motif of the LATS with threonine (T1079) required for the activation of LATS by HIPPO phosphorylation highlighted by red rectangle. **(B)** The auto-activation T-loop in LATS is 100 % conserved across the species (S909 is indicated by red rectangle). **(C)** Highly conserved motif in MOB required for regulation of MST-SAV-LATS-MOB complex and LATS auto-activation (T35 is indicated by the red rectangle. Colour scheme for amino acid residues: Blue- Hydrophobic, Red- Positive charge, Magenta- Negative charge, Green- Polar & Cyan- Aromatic. **(D)** Predicted LATS phosphorylation site in *Hvul*_YAP at S74 (homologous to the mammalian S127) and S276 (homologous to the mammalian S381). **(E)** Phosphodegron motif predicted in *Hvul*_YAP immediately downstream to the S276 (S381 in humans).

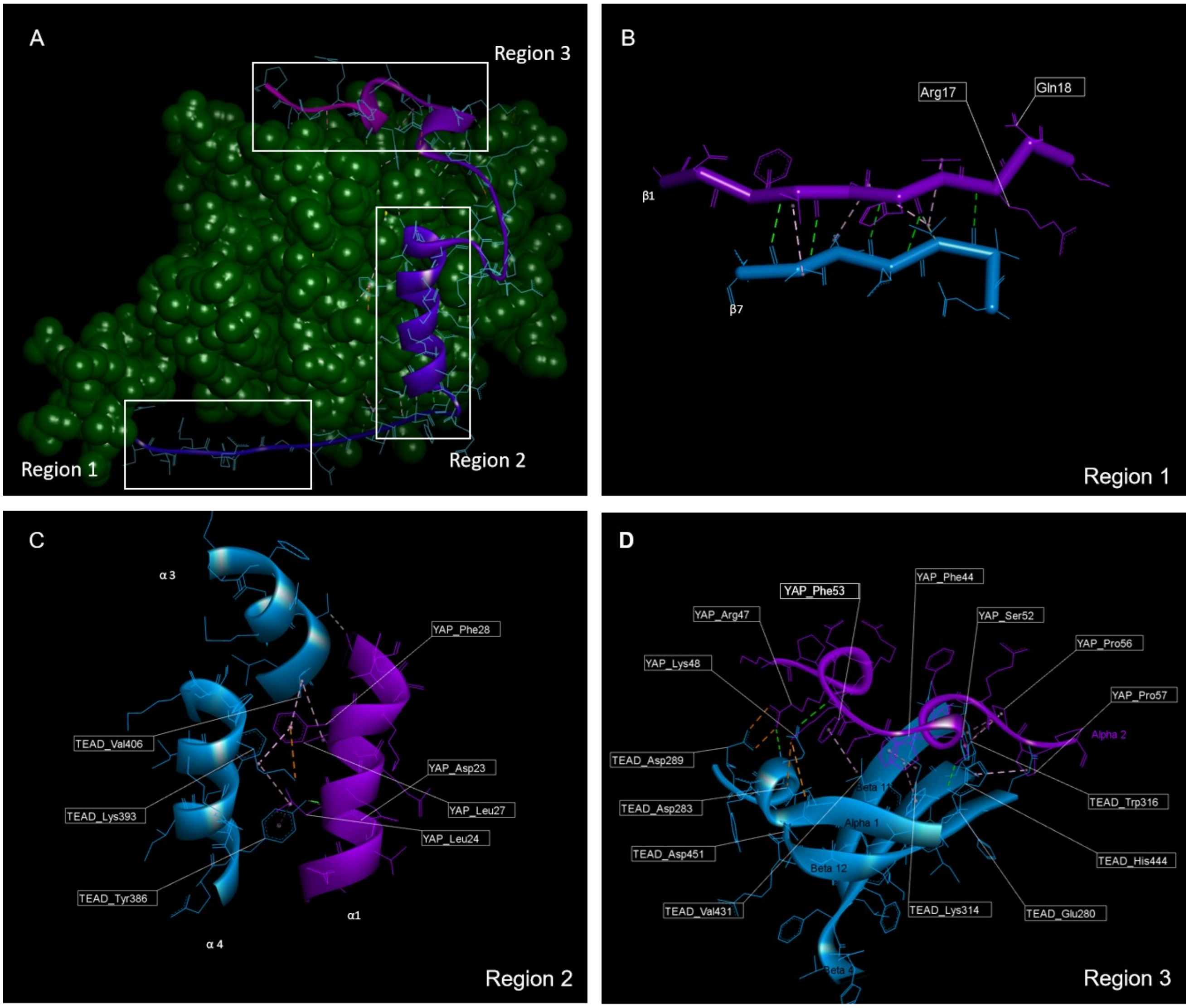


**Supplementary Figure 3. 3D structural model of YBD and TBD in the *Hydra* YAP-TEAD complex.** The 3D structure of the YBD and TBD of *Hvul*_YAP and *Hvul*_TEAD was modelled on human PDB structural model of YBD-TBD (4RE1) using MODELLER software. **(A)** The modeled YBD-TBD complex of *Hydra* showing three different regions- Region 1, Region 2 & Region 3 of TBD (purple) interacting with the globular YBD (green). **(B)** The Region 1 interface forming an anti-parallel β sheet consisting of TBD β1 and YBD β7 strands of *Hydra* interacts with only six hydrogen bonds (green dotted lines) due to the presence of Gln18 in β1 instead of Gly59 found in humans. **(C)** Region 2 has the α1 helix of the TBD (amino acids 20-32) fitting right into the binding groove of the YBD formed by the α3 and α4 helices of the YBD (amino acids 385-409). This interaction mainly consists of Leu24, Leu27and Phe28 from TBD and Try386, Lys393 and Val406 of YBD (pink dotted lines). **(D)** The region 3 in *Hydra*, has hydrophobic side chains of the TBD – Phe44 (Met86 in humans), Leu49, Pro50 and Phe53 forming extensive van der Waals interactions with the YBD of TEAD at Glu280, Ala281, Ile282, Gln286, Ile287, Leu312, Leu316, Val431, His444 & Phe446. The interface is further strengthened by multiple hydrogen bonds (indicated in green dotted lines). A predicted ability to form two salt bridges (orange dotted lines) in this region in *Hydra* model as compared to just one in the humans- TBD_Arg47:YBD_Asp289:YBD_Asp289 and TBD_Lys48:YBD_Asp283:YBD_Asp451.

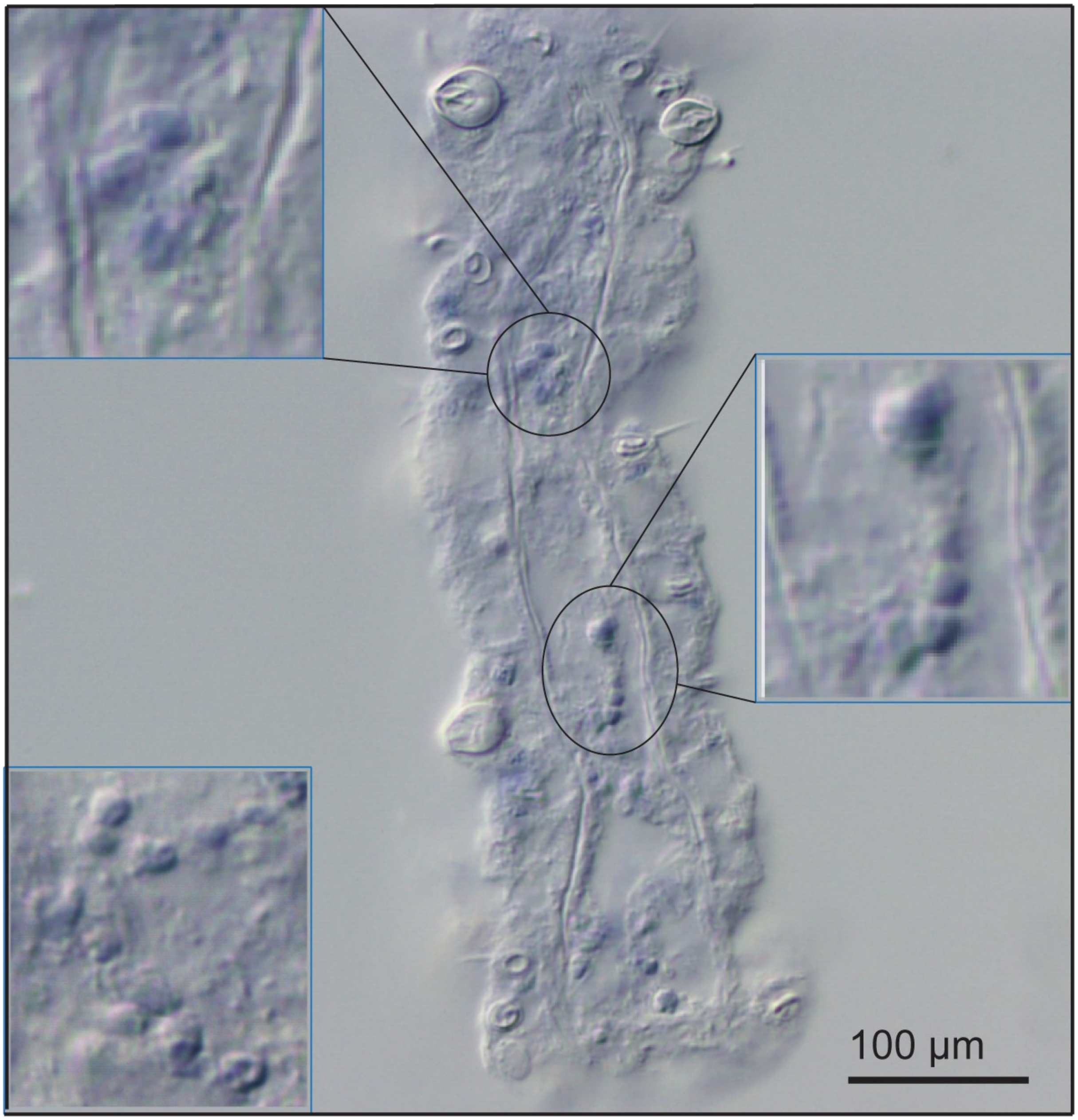


**Supplementary Figure 4. Cryosection of YAP WISH at tentacle region showing cell-type specific expression**. Crysectioning of the polyps which were stained for YAP expression by WISH were done by embedding them in PVP. These embedded polyps were cut into 25 µm sections using cryotome and then imaged. The image reveals cells in doublets, quadruplets and groups of cells among other stained cells indicating their interstitial stem cell origin, plausibly nematoblast and nests of nematoblasts. (insets shows zoomed-in areas indicted by black circles)

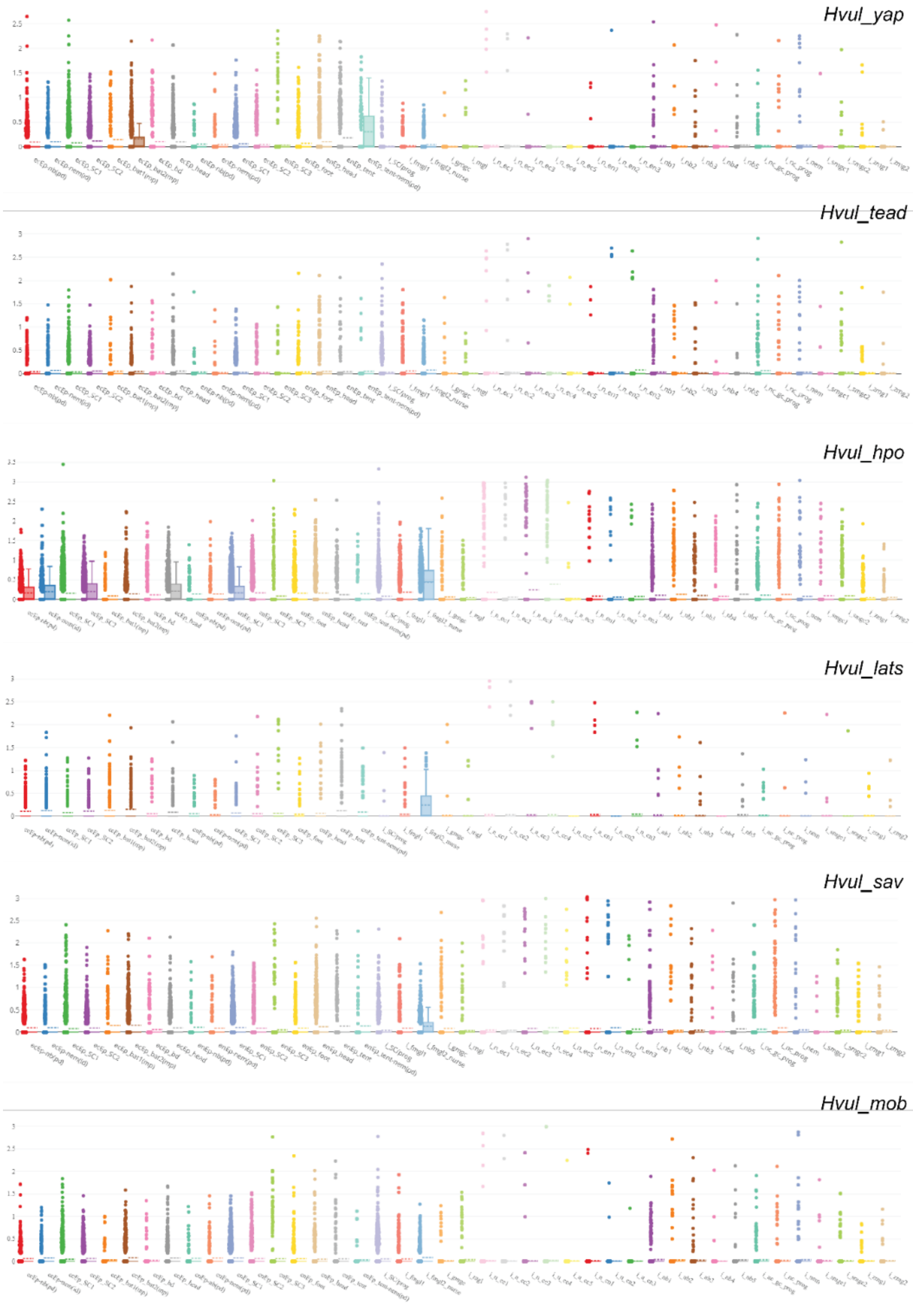


**Supplementary Figure 5. Differential expression of the Hippo pathway components in various cell types.** The differential expression of six *Hydra* Hippo genes and was examined using the Single Cell Portal. The expression pattern for Hvul_yap, Hvul_tead, Hvul_hpo, Hvul_lats, Hvul_sav & Hvul_mob is enlisted respectively from top to bottom of the figure. Cluster label abbreviation key: bat: battery cell, bd: basal disk, db: doublet cluster, ec: ectoderm, ecEP: ectodermal epithelial cell, en: endoderm, enEP: endodermal epithelial cell, fmgl: female germ-line, gc: gland cell, gmgc: granular mucous gland cell, i: cell of the interstitial lineage, id: integration doublet, mgl: male germline, mp: multiplet, nb: nematoblast, n: neuronal cell, nem: nematocyte, pd: suspected phagocytosis doublet, prog: progenitor, SC: stem cell, smgc: spumous mucous gland cell, tent: tentacle, zmg: zymogen gland cell.

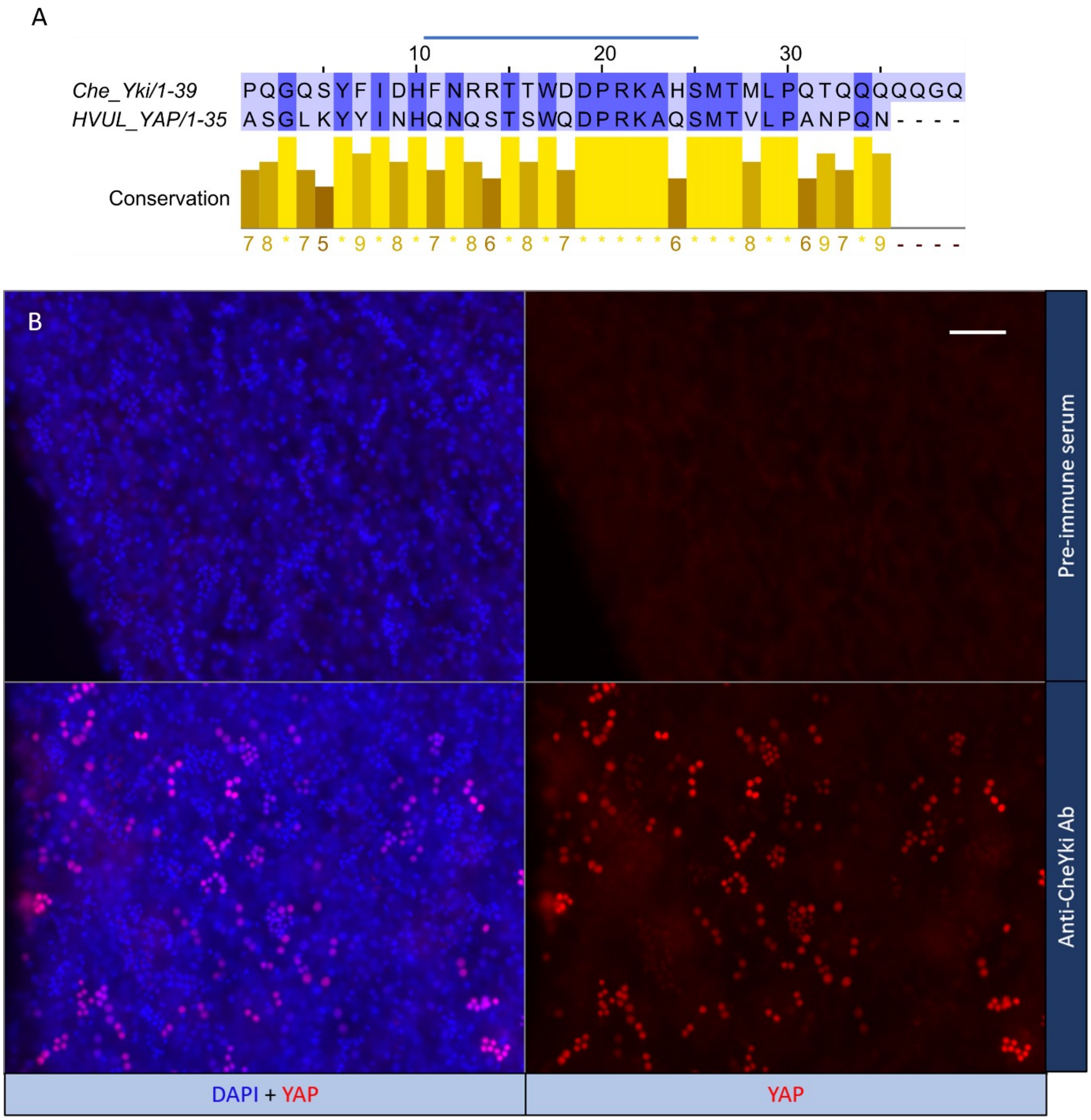


**Supplementary Figure 6. Validation of CheYki antibody in *Hydra* by immunofluorescence assay (IFA)(A)** A peptide-specific (immunogen) alignment between *Clytia* Yki and *Hydra* YAP showing a 66% similarity and a 60 % identity at motif. **(B)** IFA in *Hydra* polyps using CheYki antibody showing staining pattern of *Hvul*_YAP against the negative control (pre-immune serum). The IFA yielded a robust signal for CheYki antibody as compared to the negative control. Red: YAP & Blue: DAPI. (Scale bar = 50 µm)

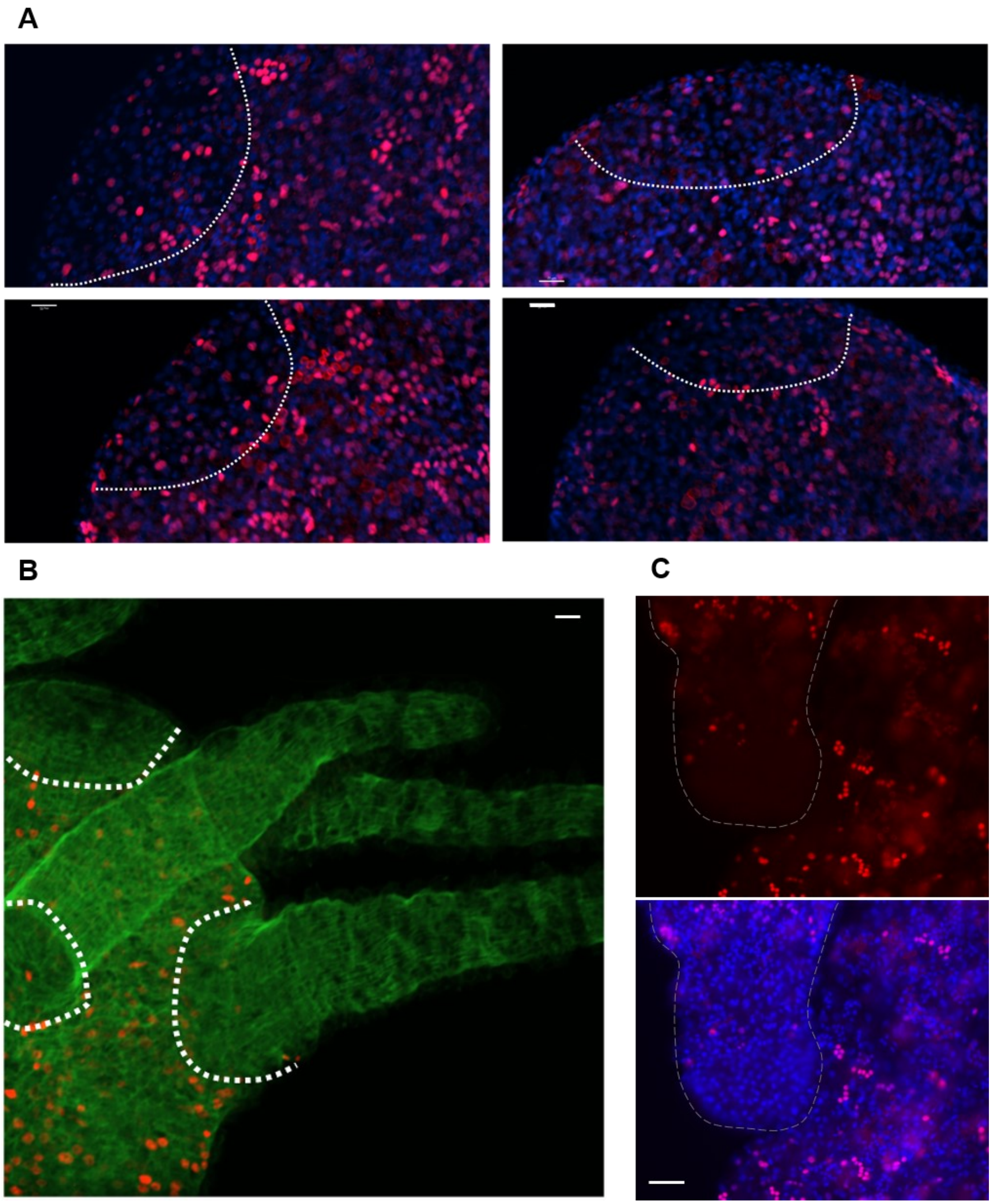


**Supplementary Figure 7.** Immunofluorescence localization of *Hvul*_YAP. The assay was performed using anti-CheYki antibody showing **(A)** YAP expressing cells at the distal tip of bud (future hypostomal region indicated by white dotted lines) non-clustered as compared to the rest of the lower bud region. Red: YAP & Blue: DAPI. (Scale bar = 50 μm) **(B)** At stage 9 onwards of budding, non-clustered YAP expressing cells were observed sparsely at the boundaries between the hypostome and the tentacle base**.** Red: YAP & Green: Actin. (Scale bar = 50 μm) **(C)** A lack of YAP expressing cells can be seen at the Adult-bud boundary (marked by dashed white line) where the future basal disk will form. The top image shows YAP expression and the bottom image shows merged image of YAP expression and nuclear stain. Red: YAP & Blue: DAPI (Scale bar = 50 μm).
